# Host traits predict the transient disease dynamics that occur during the loss of an alternative host

**DOI:** 10.64898/2026.10.28.762396

**Authors:** Miles L. Robertson, Michael H. Cortez

**Affiliations:** Department of Biological Science, Florida State University, Tallahassee, FL 32306

**Author notes:** Data accessibility statement: No new data generated in this study. Code available for download on Zenodo: https://zenodo.org/records/23775636. Statement of Authorship: MLR and MHC designed the study. MLR ran simulations and lead analysis. MLR was the lead on writing and wrote the first draft. MHC contributed to editing.

**Keywords:** transient dynamics, multi-host disease, dilution effect, amplification, alternative host traits

## Abstract

The composition of host communities in host-pathogen systems is changing through the loss and introduction of host species. Prior studies have shown that these changes can increase or decrease disease levels. However, it remains unclear what transient dynamics occur while host community composition changes, and how the traits of added or lost hosts affect those disease dynamics. In this study, we analyze a two-host model with an environmentally transmitted pathogen, where infection prevalence (proportion of infected individuals) in the focal host species changes as the second (alternative) host species is lost from the system. We use global sensitivity analysis to determine which and when alternative host traits have the strongest effects on the focal host dynamics. We find focal host infection prevalence increased or decreased monotonically when the alternative host has sufficiently higher or lower abilities to spread disease, respectively, than the focal host, but can change nonmonotonically when hosts have similar abilities to spread disease and the alternative host is a sufficiently strong competitor. Additionally, we identify that the alternative host’s shedding and uptake rates of pathogen propagules, birth rate, and interspecific competitive ability almost always have strong influence on the magnitude and rate of change in focal host infection prevalence. In contrast, The alternative host’s transmission rate, recovery rate, and disease mortality rate are only strongly influential in limited circumstances. Our study yields insight into how host traits influence the transient disease dynamics that occur following host losses in multi-host communities.

## 1 Introduction

Climate change and anthropogenic activity are causing changes in the composition of ecological communities [Travis et al., 2013], with species gains in some places and losses in others. In host-pathogen systems, the composition and biodiversity of the host community can change due to the establishment of new host species and losses of host species. These changes in host biodiversity can increase or decrease disease levels in populations and infection risk for individuals [Keesing et al., 2009, Cecere et al., 1997, Clay et al., 2009, Hall et al., 2009, Hydeman et al., 2017, Levine et al., 2017]. For example, amphibian communities in California infected with *Ribeiroia ondatrae*, a flatworm, showed a negative correlation between community disease levels and host richness [Johnson et al., 2013]. Such negative correlations between disease levels and host biodiversity are called a dilution effect [Ostfeld and Keesing, 2000]. On the other hand, avian communities in Atlanta, Georgia demonstrated a positive correlation between the community prevalence of West Nile Virus infection and host richness [Levine et al., 2017]. This positive relationship between community disease levels and host biodiveristy is called an amplification effect [Wood et al., 2014]. Meta-analyses have shown that across empirical systems, the relationships between host biodiversity and disease can be negative (dilution), positive (amplification), or nonmonotonic [Civitello et al., 2015, Halliday and Rohr, 2019].

An important factor influencing the shapes of host biodiversity-disease relationships are the disease-related and non-disease-related traits of the host species. Disease-related traits include traits like transmission and infection rates and the shedding and uptake of infectious propagules. Disease-related traits contribute to host competence, a composite trait that quantifies the ability of an exposed individual to transmit the pathogen [Stewart Merrill and Johnson, 2020, Mommer et al., 2026]. Non-disease-related traits include traits like competitive ability and defense against predation. Empirical research has shown that a host species’ traits can influence whether it will have an amplifying or diluting effect on another host species. For example, in the water flea *Daphnia dentifera*, the proportion of individuals infected with a fungal parasite (*Metschnikowia bicuspidata*) increased in the presence of the high-competence host *D. lumholtzi* [Searle et al., 2016, *Cortez, 2024]*. In contrast, the proportion of infected individuals decreased in the presence of the low-competence host D. pulicaria [Hall et al., 2009]. As another example, in some amphibian communities, Gastrophryne species consume and therefore remove zoospores of the fungus *Batrachochytrium dendrobatidis* at a much higher rate than *Bufo* species. This difference in traits explains why the presence of *Gastrophryne* species decreases infection prevalence in sympatric amphibian species, while the presence of *Bufo* species increases infection prevalence [Venesky et al., 2014]. These empirical studies together suggest that the traits of host species can affect how changes in host biodiversity affect disease levels in populations.

The importance of host traits is supported by theory [Cortez, 2021, O’Regan et al., 2015, Mihaljevic et al., 2014]. Prior theoretical studies generally predict that high competence hosts (e.g., those with high transmission and shedding rates and low mortality rates) increase disease levels when present, unless those hosts have strong competitive effects on other high-competence hosts [Cortez, 2024, Dobson, 2004, Faust et al., 2018]. Similarly, low competence hosts (e.g., those with low transmission and shedding rates and high mortality rates) decrease disease levels when present, unless those hosts have strong competitive effects on other low-competence hosts [Cortez, 2024, Rudolf and Antonovics, 2005]. Thus, the disease-related and non-disease-related traits of host species can influence how changes in host biodiversity affect disease levels in populations.

While theory shows that host traits can have important effects, our understanding of how host traits affect disease dynamics following changes in host biodiversity (e.g., the gain or loss of host species) is limited because current theory almost exclusively focuses on systems at equilibrium [e.g., Roberts and Heesterbeek, 2013, Santos et al., 2022, Bjørnstad et al., 2002, Holme and Masuda, 2015, Cortez, 2024]. Studies analyzing the basic reproduction number (*R*_0_) focus on systems near the disease-free equilibrium, while studies on the density or proportion of infected individuals in populations primarily focus on systems at endemic equilibria. Very few studies analyze host biodiversity-disease relationships for systems away from equilibrium [e.g., Roche et al., 2012, Hanthanan Arachchilage et al., 2023]. However, such information can be consequential for our understanding of disease dynamics. For example, using a two-host model, Hanthanan Arachchilage et al. [2023] found that increasing the competitive ability of an invading host increased infection prevalence in the other host on short time scales but decreased it on longer time scales. Other modeling studies have also shown that the magnitude of the diluting or amplifying effect of a host species can vary over time [Strauss et al., 2015, Hall et al., 2009]. While this prior work shows that diluting and amplifying effects differ across time scales, we lack theory that predicts how host traits influence when amplifying or diluting effects become larger or smaller through time.

Empirical work tracking disease dynamics following host addition or loss is similarly sparse, but a few studies have tracked how diluting and amplifying effects vary through time [Strauss et al., 2015, Dallas et al., 2016]. For example, Dallas et al. [2016] showed that increases in the density of *D. pulicaria* (a competing, non-susceptible host) increased the prevalence of infection by *M. bicuspidata* in *D. dentifera* on short time scales but decreased it on longer time scales during an epidemic. This demonstrates that predictions about amplifying or diluting effects at one time scale may not hold at other time scales. Altogether, this means that while we know that the gain or loss of a host species from a community can alter disease dynamics, we have a limited understanding of how those effects differ across time scales and how host trait influence those patterns.

In this study, our goal is to characterize the dynamical changes in infection prevalence of a focal host that occur as an alternative host is gradually lost from the system and to identify which alternative host traits are the most important for predicting which specific pattern occurs. To do this, we analyze a model of a two-host system with an environmentally transmitted pathogen. This community is composed of a focal and alternative host, where the alternative host is gradually lost from the system. This reflects the scenario where an environmental change has gradual impacts on host community composition, affecting some hosts negatively and others very little. We use the model to explore how loss of the alternative host affects the proportion of infected individuals (i.e., infection prevalence) in the focal host, and how those temporal patterns depend on the disease and non-disease traits of the alternative host. Specifically, we answer three questions: (i) How does focal host infection prevalence change during the loss of an alternative host? (ii) Which alternative host traits have the largest influence on whether focal host infection prevalence is higher or lower during the transient dynamics? (iii) Which alternative host traits have the largest influence on how quickly focal host infection prevalence changes? We answer these questions using local and global sensitivity analysis, which allows us to quantify the relative impact of each host trait on transient dynamics. That information allows us to determine which alternative host traits are most important for predicting and shaping the transient dynamics of the focal host.

## 2 Model

To explore how alternative host traits affect the dynamics of focal host infection prevalence as an alternative host is lost from the system, we analyze a two-host disease model with an environmentally transmitted pathogen. Our model is inspired by *Daphnia* spp. and their pathogens [Davenport et al., 2024, Stewart Merrill and Cáceres, 2018, McIntire et al., 2025, Searle et al., 2016]. For those species, susceptible individuals become infected when they encounter pathogen propagules that were released by infected individuals into the environment. The model is a system of ordinary differential equations that define the change in the densities of susceptible and infected individuals in both a focal host species (*S*_*f*_, *I*_*f*_ ) and an alternative host species (*S*_*a*_, *I*_*a*_), as well as the density of infectious propagules in the environment (*P* ). The model is

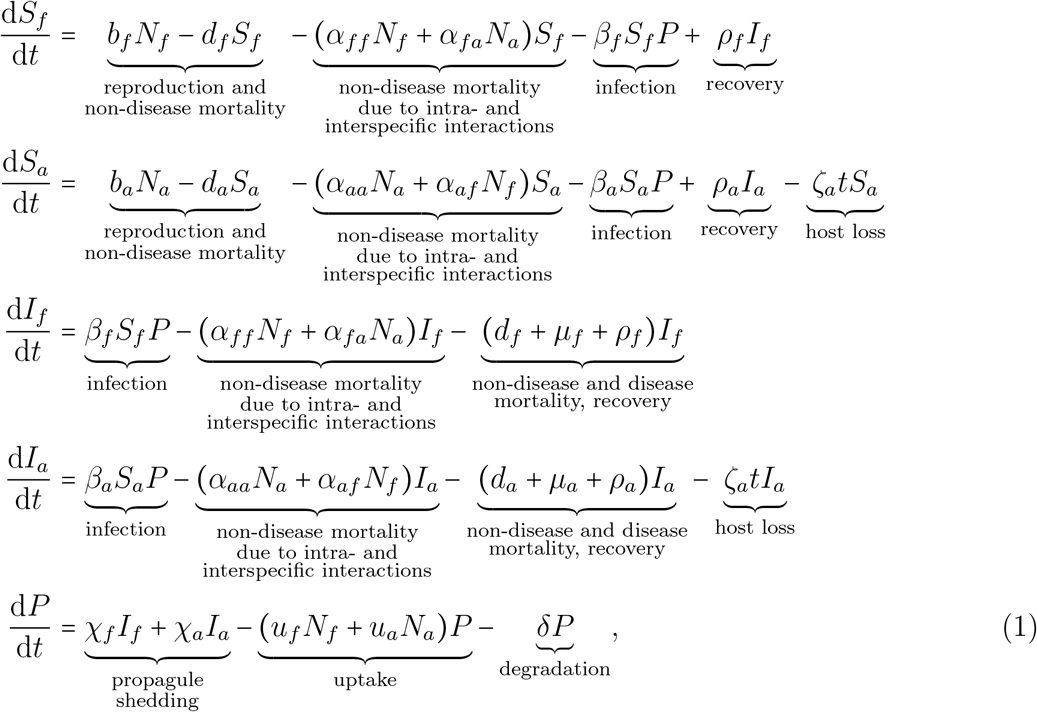

where *N*_*i*_ *S*_*i*_ *I*_*i*_ are the total host densities for each population. Susceptible density in each species increases with reproduction (*b*_*i*_*N*_*i*_); decreases due to non-disease mortality (*d*_*i*_*S*_*i*_), intraspecific and interspecific interactions ( [*α*_*ii*_*N*_*i*_ + *α*_*ij*_*N*_*j*_] *S*_*i*_), and infection (*β*_*i*_*S*_*i*_*P* ); and increases due to recovery of infected individuals (*ρ*_*i*_*I*_*i*_). The infected density for each species increases due to infection (*β*_*i*_*S*_*i*_*P* ) and decreases due to non-disease mortality due to intraspecific and interspecific interactions ( [*α*_*ii*_*N*_*i*_ + *α*_*ij*_*N*_*j*_ ]*I*_*i*_), non-disease and disease mortality ( [*d*_*i*_ + *µ*_*i*_ ]*I*_*i*_), and recovery (*ρ*_*i*_*I*_*i*_). We model loss of the alternative host using the linearly-increasing per capita rate *ζ*_*a*_*t*. Environmental propagule density increases through shedding by infected individuals (*χ*_*f*_ *I*_*f*_ + *χ*_*a*_*I*_*a*_) and decreases through uptake by all hosts ( *u*_*f*_ *N*_*f*_ + *u*_*a*_*N*_*a*_ *P* ) and degradation (*δP* ).

Our model makes the following simplifying assumptions that align with prior modeling work on *Daphnia* and other systems. First, intraspecific and interspecific host competition increase host mortality rates instead of host recruitment rates, such that per-capita host recruitment rates are density-independent and per-capita host non-disease mortality rates are density-dependent [Potapov et al., 2012, Barlow, 1996, McCormack and Allen, 2007, Roberts and Heesterbeek, 2018]. Second, we model infection rates with the commonly-used bilinear form, *βS*_*i*_*P* [e.g., Strauss et al., 2015, Searle et al., 2016, Clay et al., 2019, Cortez, 2024], which facilitates analytical calculations. Third, as in other studies [Cortez, 2024, Strauss et al., 2015, Dallas et al., 2016], we reduce model complexity by assuming that recovery does not convey any immunity to subsequent infection and recovered individuals immediately enter the susceptible class. Fourth, we assume infection status does not affect host uptake rates, birth rates, or intraspecific and interspecific interaction coefficients.

This greatly reduces the number of model parameters and is a simplifying assumption other studies also employ [e.g., Earn et al., 2000, McIntire et al., 2025]. Finally, we assume infectious propagules are shed into a shared, well-mixed environment and all propagules are identical, regardless of the host that shed them. This assumption is common in ordinary differential equation models [Searle et al., 2016, Strauss et al., 2015, Cortez, 2021].

To facilitate the analysis of the model, we convert model (1) to a form that tracks each host’s total density (*N*_*i*_ = *S*_*i*_ + *I*_*i*_) and infection prevalence (i.e., proportion of infected individuals, *Y*_*i*_ = *I*_*i*_/*N*_*i*_) of each host population. The model is

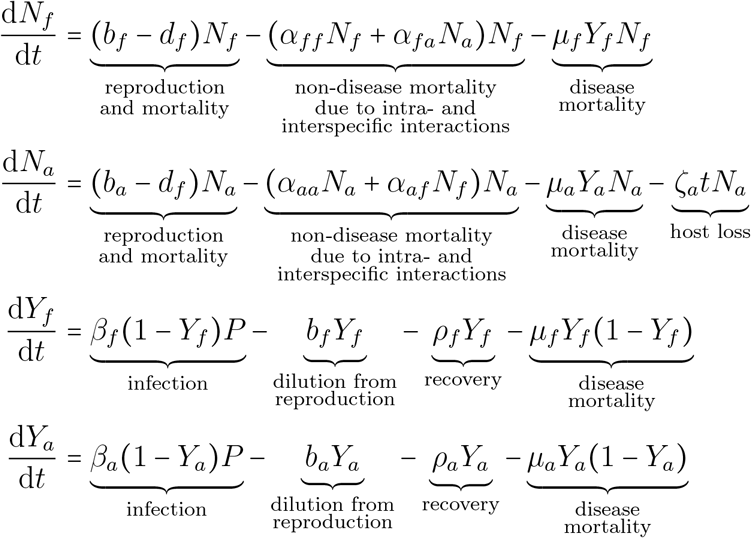

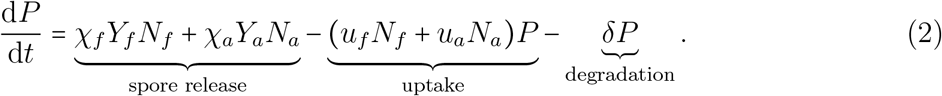

We non-dimensionalize model (2) using the following scaling:

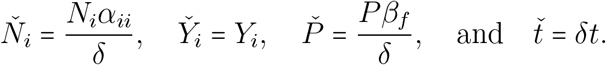

We define the following dimensionless parameters:

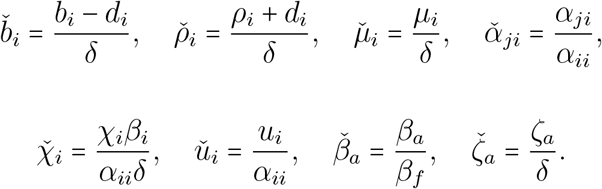

To simplify notation, we drop the wedge accent from the dimensionless parameters and variables, yielding the non-dimensionalized model. The dimensionless model is

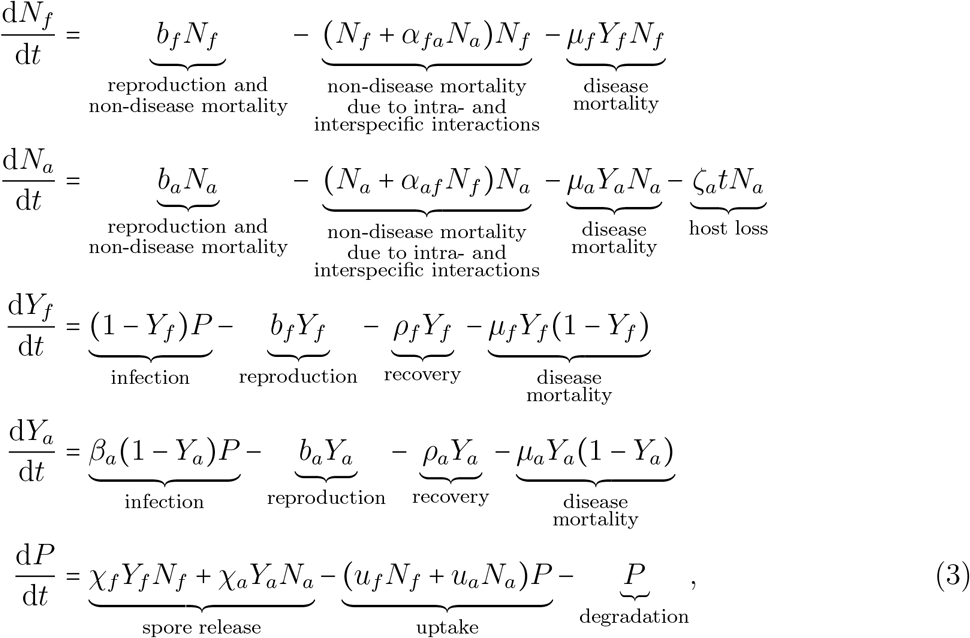

with state variables and parameters described in Table 1.

**Table 1.** State variables and parameters used in model (1).

| Symbol | Description |
| --- | --- |
| *State Variables* |  |
| $N_f$ | Focal host total density |
| $N_a$ | Alternative host total density |
| $Y_f$ | Focal host infection prevalence |
| $Y_a$ | Alternative host infection prevalence |
| $P$ | Environmental propagule density |
| $t$ | Time |
| *Pathogen Parameter* |  |
| $\delta$ | Pathogen propagule degradation rate |
| *Loss Parameter* |  |
| $\zeta_a$ | Alternative host loss parameter |
| *Disease Parameters* |  |
| $b_f$ | Focal host per capita birth rate |
| $b_a$ | Alternative host per capita birth rate |
| $d_f$ | Focal host per capita non-disease mortality rate |
| $d_a$ | Alternative host per capita non-disease mortality rate |
| $\alpha_{ff}$ | Focal host intraspecific competitive ability |
| $\alpha_{aa}$ | Alternative host intraspecific competitive ability |
| $\alpha_{af}$ | Focal host interspecific competitive ability |
| $\alpha_{fa}$ | Alternative host interspecific competitive ability |
| *Non-Disease Parameters* |  |
| $\chi_f$ | Focal host shedding rate |
| $\chi_a$ | Alternative host shedding rate |
| $u_f$ | Focal host uptake rate |
| $u_a$ | Alternative host uptake rate |
| $\beta_f$ | Focal host transmission coefficient |
| $\beta_a$ | Alternative host transmission coefficient |
| $\rho_f$ | Focal host recovery rate |
| $\rho_a$ | Alternative host recovery rate |
| $\mu_f$ | Focal host disease mortality rate |
| $\mu_a$ | Alternative host disease mortality rate |

In this study, we restrict our analysis to systems that satisfy three properties. First, the two host species must stably coexist in absence of the pathogen, which requires that interspecific host competition is less than intraspecific host competition (*α*_*fa*_*α*_*af*_ < 1).

State variables at this disease-free coexistence equilibrium are denoted with tildes, e.g.,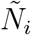. Second, the two-host endemic equilibrium is the unique attractor of the system. State variables at this two-host endemic equilibrium are denoted with asterisks, e.g., 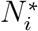. Third, the single-host endemic equilibrium for the focal host is the unique attractor in the subsystem where the alternative host is absent (see Section S1.1). State variables at this single-host endemic equilibrium are denoted with carets, e.g., 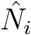.

### 2.1 Alternative Host Ability to Spread Disease

To compare the ability of each host species to spread the pathogen, we use two metrics. The first metric is the pathogen’s basic reproduction number for each host in isolation, *R*_0*i*_ [Roberts and Heesterbeek, 2013, Santos et al., 2022, Bjørnstad et al., 2002, Holme and Masuda, 2015, Cortez, 2024]. Using the Next Generation Matrix [Diekmann et al., 1990] (see Section S1.4), the value for host *i* is

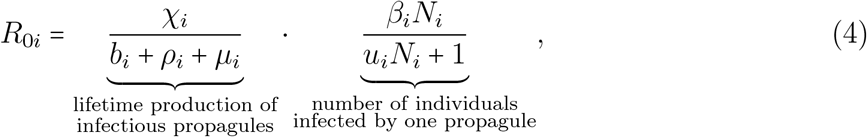

where *N*_*i*_ = *b*_*i*_ is the density of host *i* at the single-host disease-free equilibrium. *R*_0*i*_ is the product of two terms: total number of propagules that an infected individual produces before recovery or death and the average number of individuals that get infected from one propagule. Biologically, *R*_0*i*_ is the expected number of new infections caused by a single infected individual in an otherwise susceptible population.

The second metric is a host’s per-infected-individual instantaneous net production rate of infectious propagules at the two-host endemic equilibrium [Cortez, 2024]. Mathematically, this quantity is defined by

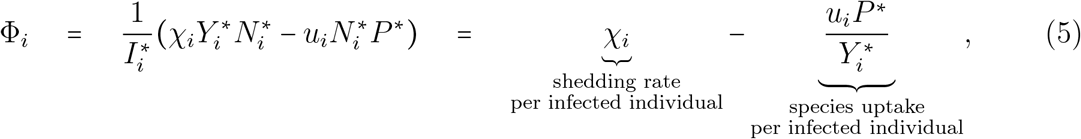

where 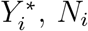 and 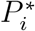 are the infection prevalence of host *i* and propagule density at the two-host endemic equilibrium, respectively. A host is a source when it is a net producer of infectious propagules, meaning its per capita shedding rate is greater than its per-infected-individual uptake rate (Φ_*i*_ 0). A host is a sink when it is a net remover of infectious propagules, meaning its per capita shedding rate is less than its per-infected-individual uptake rate (Φ_*i*_ 0).

We apply both of these metrics in different contexts. First, we use the basic reproduction number (*R*_0*i*_) to select nominal parameter sets for numerical simulation. Second, source/sink (Φ_*i*_) is a critical component of the analytical formulas we derive for local sensitivity analysis. In the discussion, we describe the connection between these metrics and host competence [Stewart Merrill and Johnson, 2020]. For all simulations in this study, all three metrics order the hosts in the same way (e.g., a host with a higher *ϕ*_*i*_ also has a higher *R*_0*i*_ and higher competence).

## 3 Study Questions and Methods

The goal of this study is to identify how alternative host traits affect the dynamical changes in focal host infection prevalence as the alternative host is lost from the system. We divide the alternative host traits in model (3) into disease traits that affect disease-related processes (shedding rate *χ*_*a*_, uptake rate *u*_*a*_, transmission coefficient *β*_*a*_, recovery rate *ρ*_*a*_, and disease mortality rate *µ*_*a*_) and non-disease traits that affect non-disease-related processes (birth rate *b*_*a*_ and interspecific competition coefficient *α*_*fa*_). We investigate three questions about how these traits influence the dynamics of focal host infection prevalence.

First, we ask, how does focal host infection prevalence change during the loss of the alternative host? Answering this question will determine when short-term and long-term changes in focal host infection prevalence are in the same or opposite directions, which yields insight into how the effects of host losses on disease depend on the time scale of interest. To answer this question, we simulate the model at nominal parameter sets spanning a broad region of parameter space. Then, we determine how the alternative host traits influence whether loss of the alternative host causes focal host infection prevalence to increase monotonically, decrease monotonically, or change nonmonotonically.

Second, we ask, which alternative host traits have the largest influence on whether focal host infection prevalence is higher or lower during the transient dynamics? Answering this question will determine which host traits are the most important for predicting the magnitude of change in focal host infection prevalence. To answer this question, we use local sensitivity analysis and global sensitivity analysis. First, we compute explicit formulas for the local sensitivity of focal host infection prevalence to each alternative host trait at the two-host endemic equilibrium. Although these formulas only necessarily apply at equilibrium, their biological interpretations can provide insight into why the signs of the effects of the alternative host traits vary across parameter space. Second, we numerically compute the global sensitivity of focal host infection prevalence to each alternative host trait. These sensitivities are computed at many time points along model simulations for a set of nominal parameter sets spanning a broad region of parameter space. This allows us to determine how the effects of the alternative host traits vary in magnitude through time and when specific traits have relatively larger versus smaller effects.

Third, we ask, which alternative host traits have the largest influence on how quickly focal host infection prevalence changes? Answering this question will determine which host traits are the most important for predicting how quickly focal host infection prevalence will change once alternative host density begins to decline. To answer this question, we use global sensitivity analysis to compute the local sensitivity of the divergence time to each alternative host trait. Here, the divergence time is the amount of time needed for focal host infection prevalence to deviate from its initial value at the two-host endemic equilibrium by 5% of the total change observed during alternative host loss. We use this metric because it indicates of how quickly focal host disease levels initially change, relative to the total amount of change.

The following provides technical details about the local sensitivity and global sensitivity analyses used to answer the second and third questions.

### 3.1 Local Sensitivity Analysis

We analytically calculate the local sensitivity of focal host infection prevalence at the two-host endemic equilibrium 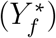 to each parameter of the alternative host (*χ*_*a*_, *u*_*a*_, *β*_*a*_, *ρ*_*a*_, *µ*_*a*_, *b*_*a*_ and *α*_*fa*_). As in our prior studies [Cortez and Duffy, 2020, Cortez, 2021, 2024], we use the method outlined in Yodzis [1988]. Let 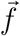 denote the system of equations in model (3) and let *J* be its Jacobian evaluated at the two-host endemic equilibrium. The sensitivity of 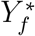 to alternative host parameter *p* is

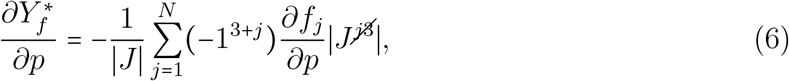

where 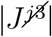 is the (*j*, 3) minor of the Jacobian, and ∣*J*∣ < 0. Positive and negative values mean focal host infection prevalence at the two-host endemic equilibrium increases and decreases, respectively, as the alternative host’s parameter value is increased.

We also compute the local sensitivity of focal host infection prevalence to a time-independent loss rate parameter. Specifically, we replace the time-dependent loss rate *ζ*_*a*_*t* in model (3) with a time-independent loss rate *η*_*a*_ and compute the local sensitivity 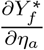. This allows us to explore how host loss affects the dynamics of focal host infection prevalence in the limit where loss of the alternative host is very slow.

### 3.2 Global Sensitivity Analysis

We applied global sensitivity analysis to nominal parameter sets spanning a broad region of parameter space. Here, we discuss (i) nominal parameter set selection, (ii) numerical simulation of model (3), (iii) our quantifications of focal host infection prevalence over time, and (iv) the global sensitivity analysis method.

#### 3.2.1 Nominal Parameter Sets

We created nominal parameter sets in the following way. First, we varied the interspecific competitive ability of each from low to high by using the *α*_*ij*_ values listed in Figure 1D. We also included the case where there is no interspecific competition (*α*_*fa*_= *α*_*af*_ = 0) as a point of comparison. Second, each host’s ability to transmit the disease was varied from low to high by using the values of *R*_0*i*_ listed in Figure 1D. These *R*_0*i*_ values were computed from particular sets of values for the 12 parameters *χ*_*i*_, *u*_*i*_, *β*_*a*_, *ρ*_*i*_ and *µ*_*i*_ (Figure 1A). These 12 individual parameters were not independently varied because it would be impractical to analyze the resulting very large number of nominal parameter sets. We used more *R*_0*a*_ values (9) than *R*_0*f*_ values (3) so that we could analyze cases with small and larger differences between the *R*_0*i*_ values without greatly increasing the number of nominal parameter sets. Third, we set the time-dependent alternative host loss parameter (*ζ*_*a*_) to 0.01.

**Figure 1.**
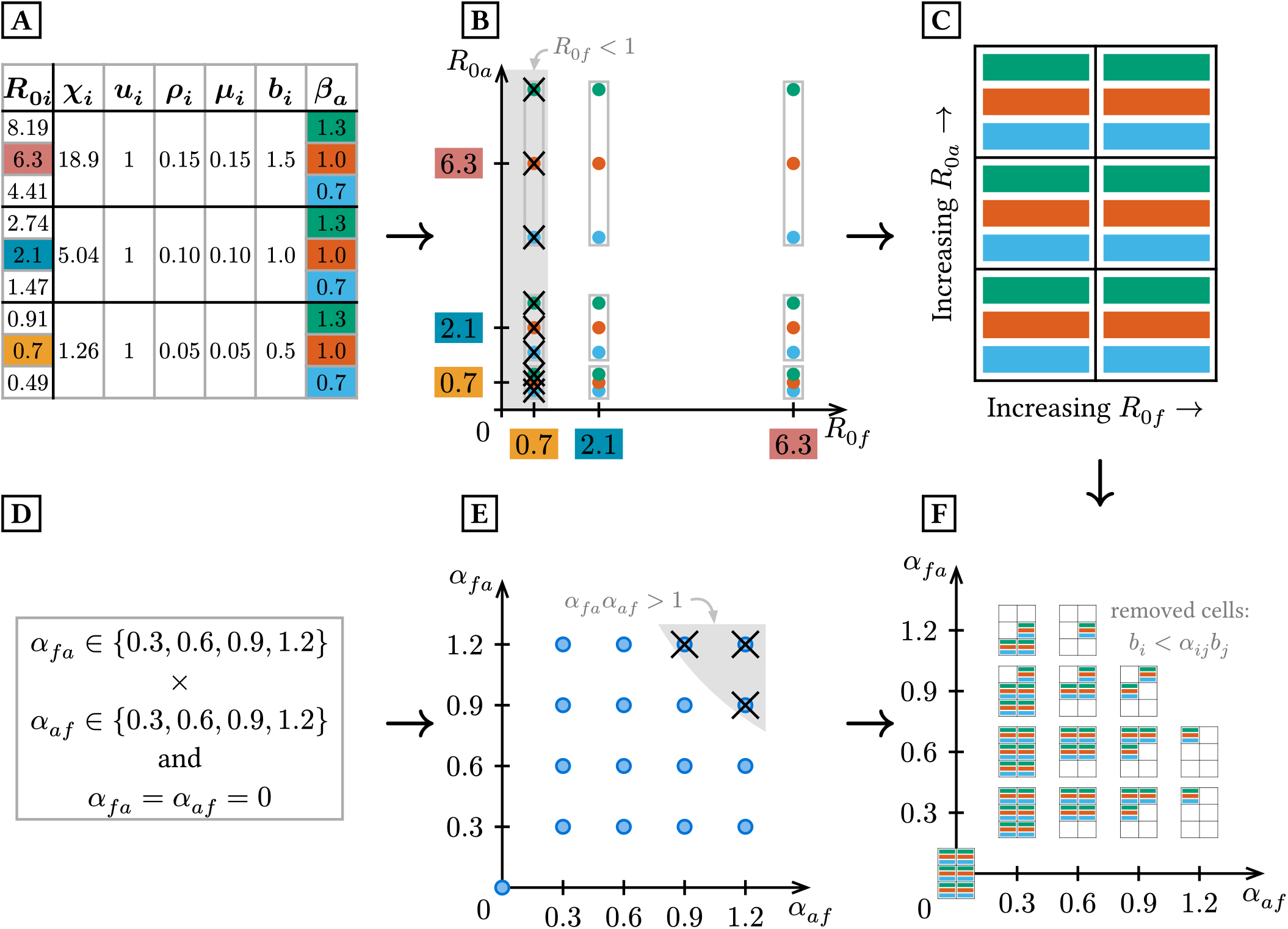
Nominal parameter set selection for model (3). **A**: Parameter values used to compute the *R*_0*i*_ values for each host, where all values of *R*_0*f*_ are computed using *β*_*f*_ = 1. **B**: Plot showing all combinations of values of *R*_0*f*_ and *R*_0*a*_, removing those where *R*_0*f*_ < 1. **C**: Simplified representation of the 18 remaining *R*_0*i*_ combinations in **B. D**: Values used for competition parameters of each host. **E**: Plot showing all combinations of values of *α*_*af*_ and *α*_*fa*_, removing those where *α*_*af*_ *α*_*fa*_ > 1. **F**: Plot showing all nominal parameter sets, generated by combinations of parameter values in **C** and **E**, removing those where *b*_*i*_ < *α*_*ij*_*b*_*j*_ for at least one host.

Finally, we factorially crossed the values of *α*_*af*_, *α*_*fa*_, *R*_0*f*_, and *R*_0*a*_, creating nominal parameter sets spanning all combinations of high to low interspecific ability (Figure 1A) and high to low *R*_0*i*_ values (Figure 1D) in each host. This yielded 459 nominal parameter sets. However, we excluded any parameter sets where (i) the focal host cannot support the disease in the absence of the alternative host (*R*_0*f*_ < 1, see Figure 1B, Section S1.1), or (ii) the two hosts cannot coexist in the absence of the disease (*α*_*fa*_*α*_*af*_ *>* 1, see Figure 1E, Section S1.2; or *b*_*i*_ < *α*_*ij*_*b*_*j*_ for either host, see Figure 1F, Section S1.2). This yields the 144 nominal parameter sets summarized in Figure 1F.

#### 3.2.2 Simulation of Model

We initialize our model starting at the two-host endemic equilibrium without alternative host loss (*t* = 0 implies *ζ*_*a*_*t* = 0), and then simulate the model through time as the alternative host is lost from the system. We used the HomotopyContinuation.jl package [Breiding and Timme, 2018] of the Julia programming language [Bezanson et al., 2017] to find the two-host endemic equilibrium. We simulated the model using Tsit5 [Tsitouras, 2011], an optimized version of RK5(4) available in the DifferentialEquations.jl package [Rackauckas and Nie, 2017]. If the alternative host density fell below 10^−10^, it was considered extinct and its density was set to zero. All simulations ran for 500 units of time, which was sufficient for the alternative host to go extinct and the system to converge to the single-host endemic equilibrium for the focal host.

#### 3.2.3 Quantities of Interest in Global Sensitivity Analysis

For question 2, the quantity of interest is focal host infection prevalence (*Y*_*f*_, Figure 2, time series values in first column). We analyzed focal host infection prevalence at 25 points in time, which were selected by rounding 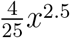 for *x* = 1…25 to the nearest integer. This sequence allowed for smaller spacing between time points during the early transient period and larger spacing later in time when the system was close to the single-host endemic equilibrium.

**Figure 2.**
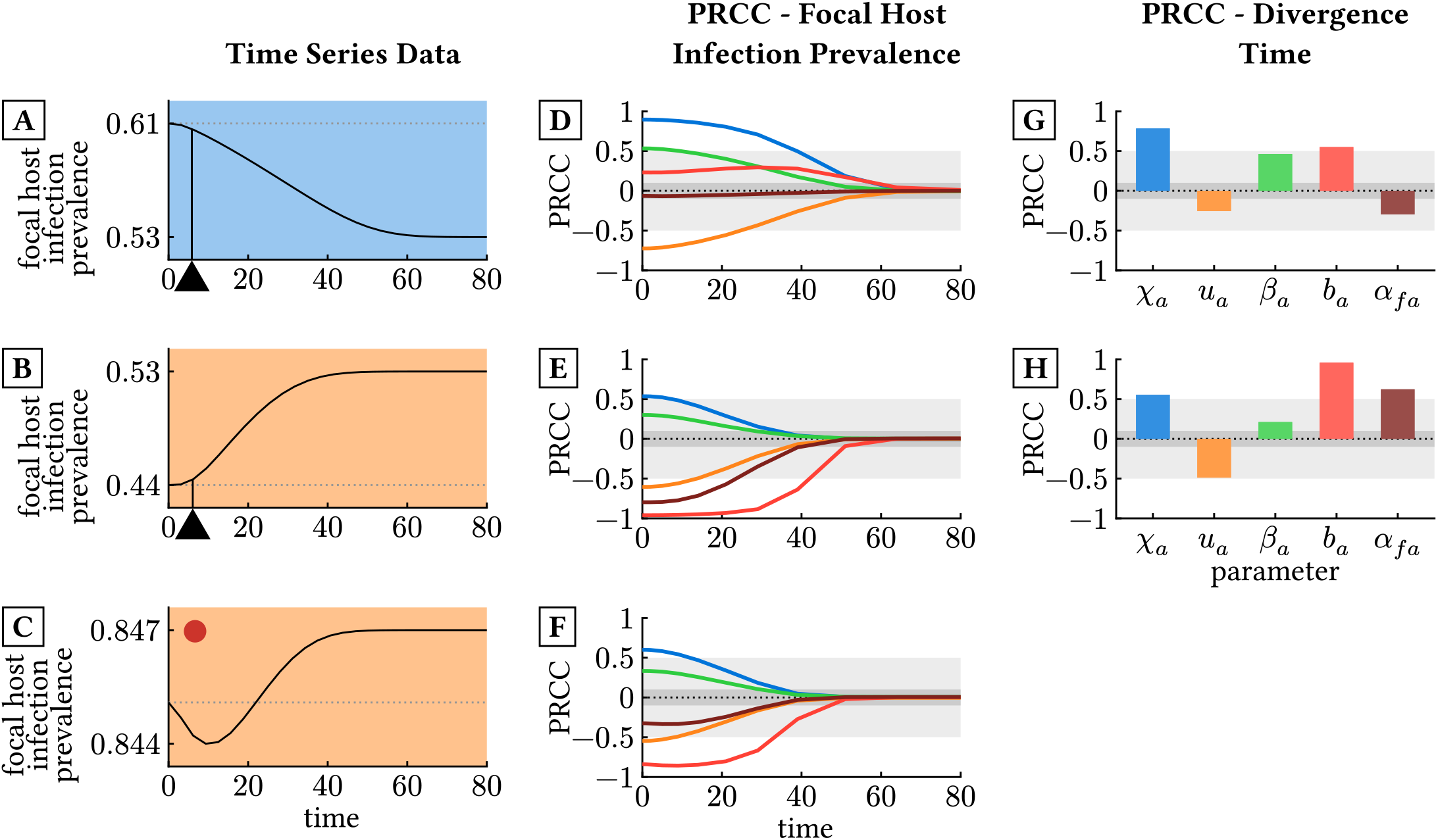
Example time series and global sensitivity results (PRCC) for three representative cases. Each row represents outcomes for a single nominal parameter set. **(A-C)** Time series of focal host infection prevalence. The blue and orange backgrounds indicate a long-term decrease or increase in focal host infection prevalence, respectively, while a red dot indicates nonmonotonic change over time. Black triangles indicate the divergence time, i.e., the first time point when focal host infection prevalence has deviated from its initial value by 5% of the total change observed over the whole simulation. **(D-** (∣PRCC∣ ≥ 0.5) and gray regions indicate when parameters are moderately influential **F)** PRCC values for focal host infection prevalence; colors correspond to parameters in panels G and H. White regions indicate where parameters are strongly influential sets are as follows. **(A**,**D**,**G)**: *α*_*af*_ = 0.6, *α*_*fa*_ = 0.3, *R*_0*f*_ = 2.1, *R*_0*a*_ = 2.74. **(B**,**E**,**H)**: indicate where parameters are strongly influential (∣PRCC∣ ≥ 0.5) and gray regions indicate or noninfluential (∣PRCC∣ < 0.5). **(G**,**H)** PRCC values for divergence time. White regions when parameters are moderately influential or noninfluential (∣PRCC∣ < 0.5). Parameter *α*_*af*_ = 0.3, *α*_*fa*_ = 0.6, *R*_0*f*_ = 6.3, *R*_0*a*_ = 0.7. **(C**,**F)**: *α*_*af*_ = 0.9, *α*_*fa*_ = 0.9, *R*_0*f*_ = 6.3, *R*_0*a*_ = 4.41.

For question 3, the quantity of interest is the divergence time (Figure 2, black arrows in first column). The divergence time is the amount of time needed for focal host infection prevalence to deviate from its initial value at the two-host endemic equilibrium by 5% of the total change observed during alternative host loss. Mathematically, the divergence time is defined by 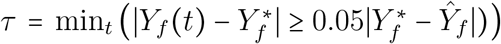. This metric was only analyzed at nominal parameter sets where focal host infection prevalence changed monotonically because the divergence time is not a continuous function of the model parameters when focal host infection prevalence changes nonmonotonically (no black arrow for row C in first column in Figure 2).

#### 3.2.4. Global Sensitivity Analysis Method: Partial Rank Correlation Coefficient

Following Hanthanan Arachchilage et al. [2023], we computed the global sensitivities of focal host infection prevalence to model parameters using Partial Rank Correlation Coefficients (PRCC) [McKay et al., 1976]. To do this, we first generated parameter sets around each nominal parameter set using Latin Hypercube Sampling (LHS) [McKay et al., 1976]. We created hypercubes around each nominal parameter set by varying each of the 14 nominal parameter values by ±10%. To satisfy the assumptions of PRCC, we verified that all quantities of interest change monotonically across each parameter range. When a parameter’s range included nonmonotonic changes in one or more quantities of interest, the parameter’s range was truncated to the monotonic region. Using a bin size of 1,000, we repeatedly generated 1,000 parameter sets from the hypercube until we collected 50,000 sets. While generating each set of 1,000 parameter sets, we only retained those that (1) yielded a unique and stable two-host endemic equilibrium, (2) satisfied *R*_0*f*_ > 1, meaning the focal host can support the disease in the absence of the alternative host, and (3) allowed for host coexistence in the absence of the pathogen (*α*_*fa*_*α*_*af*_ < 1, *b*_*f*_ > *α*_*fa*_*b*_*a*_, and *b*_*a*_ > *α*_*af*_ *b*_*f*_ ). This first step of the method was slightly altered for nominal parameter sets where interspecific competition was absent (*α*_*fa*_ = *α*_*af*_ = 0). At these nominal parameter sets, we did not vary the *α*_*ij*_ values; this avoided using one-sided intervals or using negative *α*_*ij*_ values, which would result in mutualistic or consumer-resource interactions between the host species.

Second, we computed PRCC values for each quantity of interest and host parameter from the 50,000 parameter sets from each hypercube. Although we computed PRCC values for all 14 parameters, our study only focuses on the 7 alternative host parameters. PRCC values range from -1 to 1. Positive and negative PRCC values indicate that an increase in the parameter increases and decreases the quantity of interest, respectively, while values closer to zero indicate weaker influence of the parameter on the quantity of interest. We categorized parameters as strongly influential when PRCC ∣ ≥0.5, moderately influential when 0.5 > ∣PRCC∣ ≥ 0.1, and noninfluential when PRCC < 0.1. In the main text, we focus on categorizing the regions of parameter space where parameters are strongly influential for at least one time point (curves and bars outside of gray region in middle and right columns of Figure 2). We provide detailed results about moderately influential and noninfluential parameters in Appendix Section S5.

## 4 Results

### 4.1 Question 1. How does focal host infection prevalence change during the loss of an alternative host?

#### 4.1.1 Local Sensitivity to Alternative Host Loss Rate (*η*_*a*_)

To assess how loss of the alternative host affects focal host infection prevalence when the loss rate is very slow, we replace the time-dependent loss rate (*ζ*_*a*_*t*) in model (3) with a time-independent loss rate (*η*_*a*_) and evaluate the sensitivity of focal host infection prevalence at equilibrium to *η*_*a*_.

The local sensitivity of focal host infection prevalence 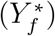 to the alternative host loss rate (*η*_*a*_) is

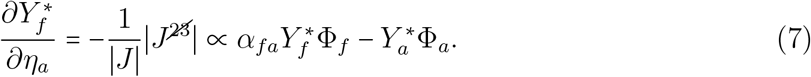

This sensitivity can have either sign. Slow loss of the alternative host increases (positive values) or decreases (negative values) focal host infection prevalence when the alternative host is a sufficiently large sink (Φ < 0) or a sufficiently source source (Φ > 0), respectively. However, if the alternative host is a sufficiently strong interspecific competitor (*α*_*fa*_ large), then the response can be reversed when the focal host is a sufficiently large source (Φ_*f*_ > 0) or sink (Φ_*f*_ < 0), respectively.

#### 4.1.2 Numerical Simulation of Nominal Parameter Sets

Across all nominal parameter sets, we find that focal host infection prevalence either (i) monotonically decreases, (ii) monotonically increases, or (iii) transiently decreases and then asymptotically increases (hereafter referred to as a nonmonotonic response). We discuss the conditions where each of these cases occurs.

First, time series that monotonically decrease (representative example in Figure 3A) only occur when *R*_0*a*_ is sufficiently high relative to *R*_0*f*_ . This threshold increases as alternative host competitive ability increases (all blue cells without red circles in top left blocks of each 3-by-2 grid in Figure 3F, with fewer such cells for lower *R*_0*a*_ values as *α*_*fa*_ increases).

**Figure 3.**
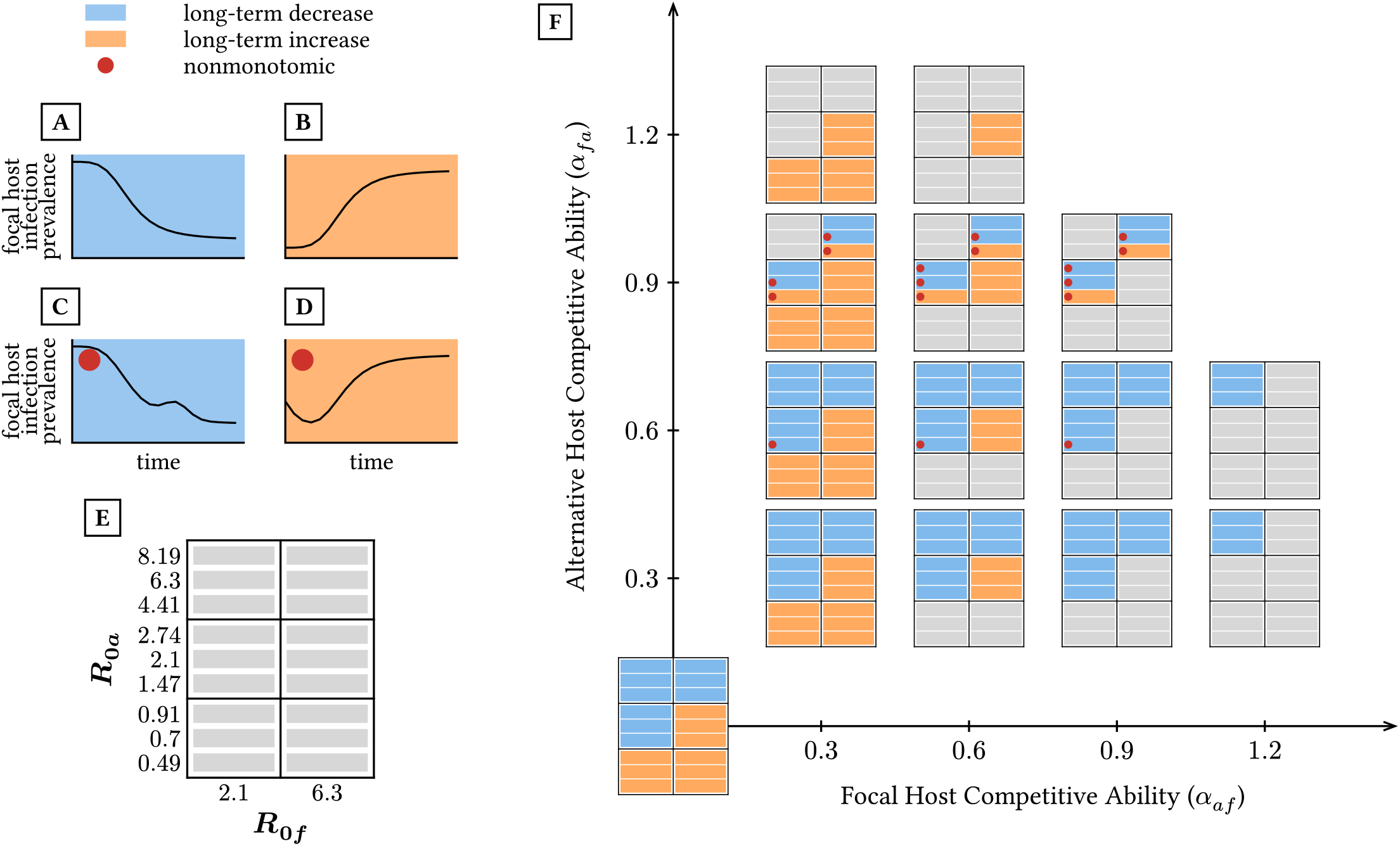
Summary of the dynamics of focal host infection prevalence at each nominal parameter set. **(A**,**B**,**C**,**D)** Representative examples of (A) monotonic decreases, (B) monotonic increases, and (C,D) nonmonotonic changes in focal host infection prevalence as the alternative host is lost from the system. **(E)** Graphical representation of host *R*_0*i*_ values for each parameter set as shown in inner plots of **F. (F)** Observed responses in focal host infection prevalence to the loss of the alternative host at each nominal parameter set. Blue and orange cells indicate that focal host infection prevalence decreased and increased, respectively, in the long term. Red dots indicate nonmonotonic changes in focal host infection prevalence. Gray cells represent nominal parameter sets that were excluded, as shown in Figure 1.

Second, time series that monotonically increase (representative example in Figure 3B) only occur when *R*_0*a*_ is significantly less than *R*_0*f*_, regardless of alternative host competitive ability (all orange cells without red circles in bottom right in Figure 3F).

Finally, time series with a nonmonotonic response only occur when *R*_0*a*_ is similar to *R*_0*f*_ and alternative host competitive ability is sufficiently high (cells with red dots occur where the values of *R*_0*i*_ are similar and when *α*_*fa*_ > 0.6 in Figure 3F). There are two kinds of nonmonotonic responses. First, focal host infection prevalence has an initial decrease followed by at least one period of increase, but the asymptotic value is never greater than the initial value (representative example in Figure 3C). Second, focal host infection prevalence transiently decreases but then increases to a value above its initial value. This case only occurs when the alternative host is a sufficiently strong competitor and when *R*_0*a*_ is slightly smaller than *R*_0*f*_ (representative example in Figure 3D).

#### 4.1.3 Summary of Question 1 Findings

The local sensitivity formula (7) show that source and sink alternative hosts typically decrease and increase focal host infection prevalence, respectively, but these can be reversed if the alternative host is a sufficiently strong competitor. Consistent with these findings, our numerical simulations show that focal host infection prevalence decreases and increases monotonically when the alternative host has sufficiently higher and lower *R*_0*i*_ value, respectively, than the focal host. However, we also found cases with nonmonotonic responses when *R*_0*i*_ values are similar and alternative host competitive ability was sufficiently high. These nonmonotonic cases are empirically important because they imply that the observed effects of alternative host loss on short time scales (e.g., short-term decrease of focal host infection prevalence) incorrectly predict the long-term effects of alternative host loss (e.g., long-term increase exceeding initial values of focal host infection prevalence).

### 4.2 Question 2. Which alternative host traits have the largest influence on whether focal host infection prevalence is higher or lower during the transient dynamics?

Here, we use local and global sensitivity analysis to determine how the alternative host traits affect focal host infection prevalence. Our explicit formulas for the local sensitivities determine the range of possible effects each alternative host parameter can have on focal host infection prevalence at the two-host endemic equilibrium. Our numerically-computed PRCC values determine the effects that alternative host traits have during transient dynamics. PRCC values range from -1 to 1, with positive and negative values indicating that an increase in the parameter increases and decreases focal host infection prevalence, respectively, while values closer to zero indicate weaker influence of the parameter on focal host infection prevalence.

For each alternative host parameter, we focus on identifying regions of parameter space where ∣PRCC ∣ > 0.5. Biologically, this corresponds to identifying if and when the alternative host trait has a strong influence on the dynamics of focal host infection prevalence. For completeness, Appendix Section S5 gives additional information about the effects of each parameter when those effects are moderately influential (0.5 ≥ ∣PRCC ∣ > 0.1) and noninfluential (∣PRCC∣ < 0.1).

Figure 4AC summarizes the regions of parameter space where each parameter is strongly influential ( ∣ PRCC∣ > 0.5). In Figure 4A, the encircled regions indicate the levels of competition for which a parameter is strongly influential for at least one combination of *R*_0*a*_ and *R*_0*f*_ values. In Figure 4C, the encircled regions indicate *R*_0*i*_ values for which a parameter is strongly influential for at least one combination of the interspecific competition coefficients (*α*_*af*_, *α*_*fa*_).

**Figure 4.**
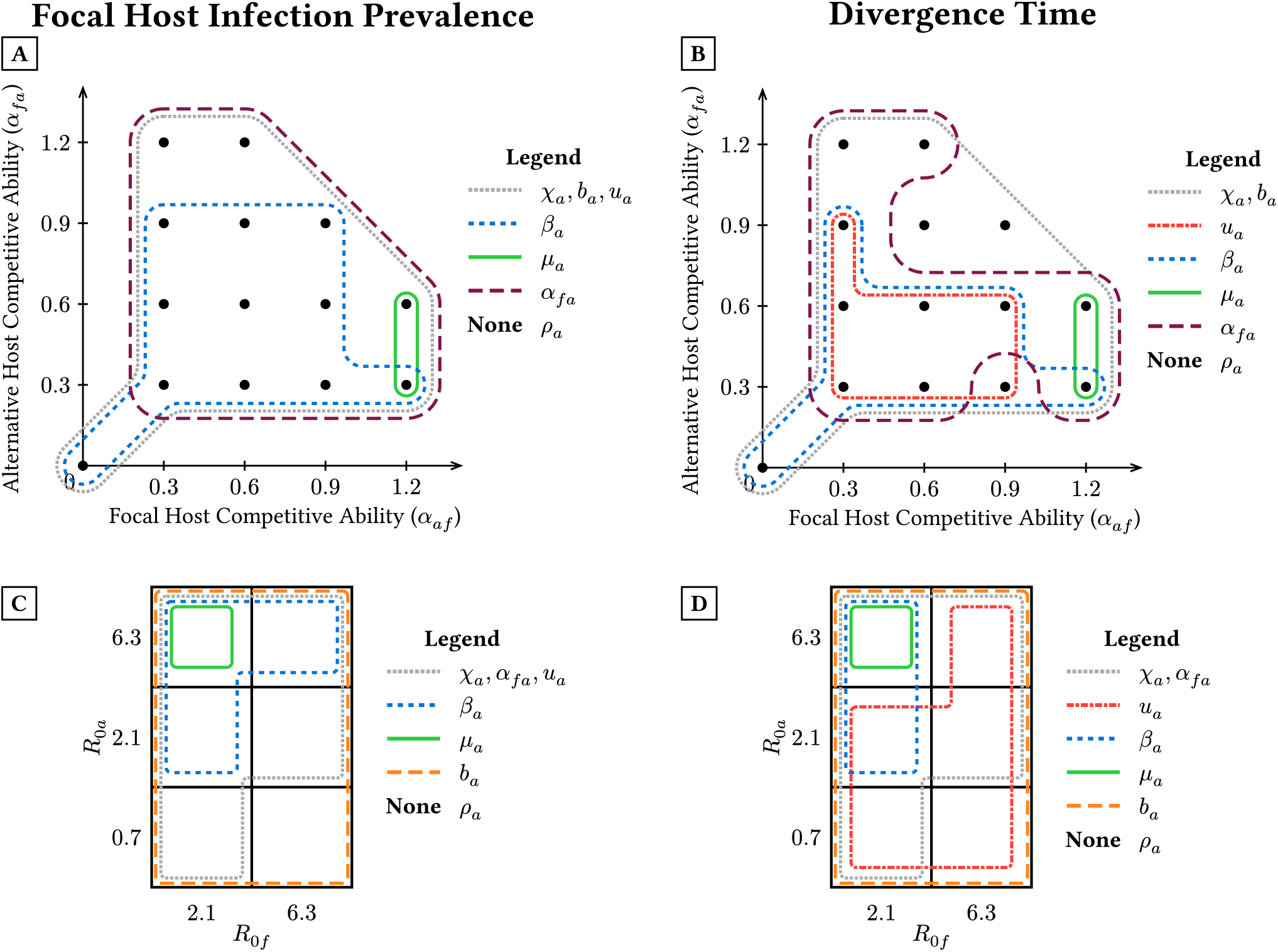
Regions of parameter space where parameters are strongly influential (∣ PRCC∣ ≥ 0.5). **(A**,**B)** Regions indicate where each alternative host parameter is strongly influential for at least one combination of *R*_0*i*_ values. **(C**,**D)** Regions indicate where each alternative host parameter is strongly influential for at least one combination of *α*_*af*_ and *α*_*fa*_values. **(A**,**C)** Regions indicate where each alternative host parameter is strongly influential for focal host infection prevalence for at least one time point. **(B**,**D)** Regions indicate where each alternative host parameter is strongly influential for the divergence time, i.e., the first time point when focal host infection prevalence has deviated from its initial value by 5% of the total change observed over the whole simulation.

#### 4.2.1 Effect of Alternative Host Shedding Rate (*χ*_*a*_) on Focal Host Infection Prevalence

The local sensitivity of focal host infection prevalence 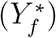 to the alternative host shedding rate (*χ*_*a*_) is

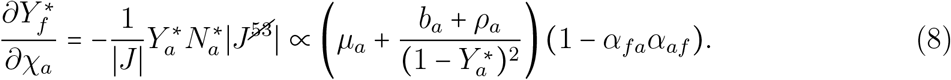

The sensitivity is positive unless *α*_*fa*_*α*_*af*_ 1. This means that an increase in alternative host shedding rate increases focal host infection prevalence at equilibrium, unless interspecific competition is sufficiently high such that the two host species cannot coexist in the absence of the pathogen (*α*_*fa*_*α*_*af*_ > 1). For all nominal parameter sets, this sensitivity is positive because we only consider systems where the hosts can stably coexist in the absence of the pathogen.

In line with local sensitivity results, the global sensitivities show that higher alternative host shedding rates increase focal host infection prevalence at all time points (see Appendix Section S5.1). These effects are strongly influential ( ∣ PRCC ∣ ≥0.5) across parameter space except when *R*_0*a*_ is less than 1 and *R*_0*f*_ is large (dotted region in Figure 4AC).

#### 4.2.2 Effect of Alternative Host Uptake Rate (*u*_*a*_) on Focal Host Infection Prevalence

The local sensitivity of focal host infection prevalence 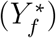 to the alternative host uptake rate (*u*_*a*_) is

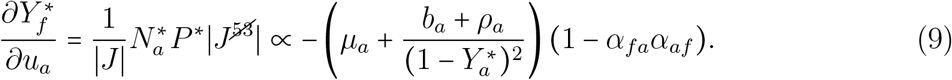

The sensitivity is negative unless *α*_*fa*_*α*_*af*_ > 1. This means that an increase in alternative host uptake rate (*u*_*a*_) decreases focal host infection prevalence at equilibrium, unless interspecific competition is sufficiently high such that the host species cannot coexist in the absence of the pathogen (*α*_*fa*_*α*_*af*_ > 1). The conditions defining the sign of this sensitivity are exactly the opposite of those for *χ*_*a*_, as the two sensitivities always have the opposite sign (see (8) and (9)). For all nominal parameter sets, this sensitivity is negative because we only consider systems where the hosts can stably coexist in the absence of the pathogen.

In line with local sensitivity results, the global sensitivities show that higher alternative host uptake rates always decrease focal host infection prevalence at all time points (see Appendix Section S5.2). These effects are strongly influential ( ∣PRCC∣ ≥ 0.5) across parameter space except when *R*_0*a*_ is less than 1 and *R*_0*f*_ is large (dotted region in Figure 4AC).

#### 4.2.3 Effect of Alternative Host Transmission Coefficient (*β*_*a*_) on Focal Host Infection Prevalence

The local sensitivity of focal host infection prevalence 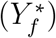 to the alternative host transmission coefficient (*β*_*a*_) is

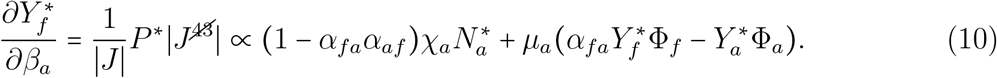

The sensitivity is positive in much of parameter space, meaning that focal host infection prevalence at equilibrium is higher when the alternative host has a higher transmission coefficient. However, the sensitivity can be negative when (i) interspecific competition is sufficiently strong such that the pathogen mediates coexistence of the hosts (*α*_*fa*_*α*_*af*_ > 1), (ii) the alternative host is a sufficiently large source (Φ_*a*_ positive and large), or (iii) the alternative host is a sufficiently strong interspecific competitor and the focal host is a sufficiently large sink (*α*_*fa*_ large and Φ_*f*_ negative and large).

The global sensitivities show that higher alternative host transmission coefficients increase focal host infection prevalence at all time points (see Appendix Section S5.3). This effect is strongly influential (∣ PRCC ∣ ≥ 0.5) when *R*_0*a*_ is similar to or greater than *R*_0*f*_ and the alternative host competitive ability is not too high (dashed blue regions in Figure 4AC). We note that the constant sign of the PRCC value is at odds with the local sensitivity formula that predicts either sign is possible; we return to this point in the Discussion.

#### 4.2.4 Effect of Alternative Host Recovery Rate (*ρ*_*a*_) on Focal Host Infection Prevalence

The local sensitivity of focal host infection prevalence 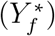 to the alternative host recovery rate (*ρ*_*a*_) is

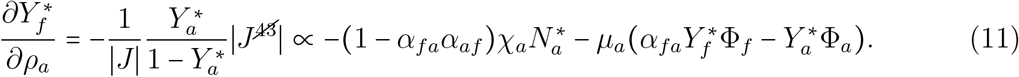

The sensitivity is negative in much of parameter space, meaning that focal host infection prevalence at equilibrium is lower when the alternative host has a higher recovery rate. However, the sensitivity can be positive when (i) interspecific competition is sufficiently strong such that the pathogen mediates coexistence of the hosts (*α*_*fa*_*α*_*af*_ 1), (ii) the alternative host is a sufficiently large source (Φ_*a*_ positive and large), or (iii) the alternative host is a sufficiently strong interspecific competitor and the focal host is a sufficiently large sink (*α*_*fa*_ large and Φ_*f*_ negative and large). These conditions defining exceptions are identical to those for *β*_*a*_, as the two sensitivities always have the opposite sign (see (10) and (11)).

The global sensitivities show that higher alternative host recovery rates decrease focal host infection prevalence at all time points (see Appendix Section S5.4). These effects are never strongly influential ( ∣PRCC ∣ ≥0.5; *ρ*_*a*_ has no regions in Figure 4AC). We note that the constant sign of the PRCC value is at odds with the local sensitivity formula that predicts either sign is possible; we return to this point in the Discussion.

#### 4.2.5 Effect of Alternative Host Disease Mortality Rate (*µ*_*a*_) on Focal Host Infection Prevalence

The local sensitivity of focal host infection prevalence 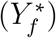 to the alternative host disease mortality rate (*µ*_*a*_) is

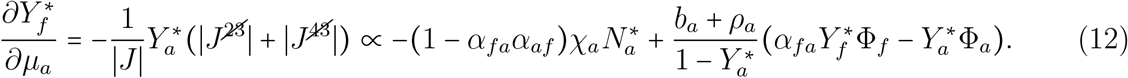

The sensitivity is negative in much of parameter space, meaning that focal host infection prevalence at equilibrium is lower when the alternative host has a higher disease mortality rate. However, the sensitivity can be positive when (i) interspecific competition is sufficiently strong such that the pathogen mediates coexistence of the hosts (*α*_*fa*_*α*_*af*_ 1), (ii) the alternative host is a sufficiently large sink (Φ_*a*_ negative and large), or (iii) the alternative host is a sufficiently strong interspecific competitor and the focal host is a sufficiently large source (*α*_*fa*_ large and Φ_*f*_ positive and large).

The global sensitivities show that higher alternative host disease mortality rates decrease focal host infection prevalence at all time points when *R*_0*a*_ is greater than or similar to *R*_0*f*_ (see Appendix Section S5.5). These effects are only strongly influential ( ∣PRCC ∣ ≥ 0.5) when *R*_0*a*_ is much greater than *R*_0*f*_ and the focal host is a strong competitor (*α*_*af*_ = 1.2; solid green region in Figure 4AC). We note that higher alternative host disease mortality rates increase focal host infection prevalence at all time points when *R*_0*a*_ is much less than *R*_0*f*_, but this effect is never strongly influential (see Appendix Section S5.5).

#### 4.2.6 Effect of Alternative Host Birth Rate (*b*_*a*_) on Focal Host Infection Prevalence

The local sensitivity of focal host infection prevalence 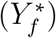 to the alternative host birth rate (*b*_*a*_) is

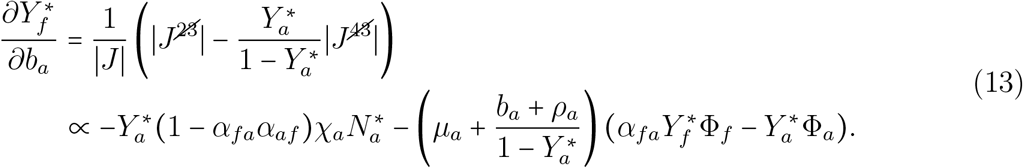

The sensitivity is negative in much of parameter space, meaning that focal host infection prevalence at equilibrium is lower when the alternative host has a higher birth rate. However, the sensitivity can be positive when (i) interspecific competition is sufficiently strong such that the pathogen mediates coexistence of the hosts (*α*_*fa*_*α*_*af*_ > 1), (ii) the alternative host is a sufficiently large source (Φ_*a*_ positive and large), or (iii) the alternative host is a sufficiently strong interspecific competitor and the focal host is a sufficiently large sink (*α*_*fa*_ large and Φ_*f*_ negative and large).

The global sensitivities show that higher alternative host birth rates almost always increase focal host infection prevalence at all time points when *R*_0*a*_ is greater than or similar to *R*_0*f*_, and decrease it when *R*_0*a*_ is much less than *R*_0*f*_ . Exceptions to this pattern only occur at or near nonmonotonic cases, where higher alternative host birth rates decrease focal host infection prevalence despite hosts having similar *R*_0*i*_ values and in one instance where higher alternative host birth rates decrease focal host infection prevalence initially, but later increase it (see Appendix Section S5.6). Across parameter space, these effects are strongly influential ( ∣PRCC ∣ ≥ 0.5; dotted black in Figure 4A and dashed orange in Figure 4C). However, there are some idiosyncratic exceptions when the host species have similar *R*_0*i*_ values (see Appendix Section S5.6 for details).

#### 4.2.7 Effect of Alternative Host Competitive Ability (*α*_*fa*_) on Focal Host Infection Prevalence

The local sensitivity of focal host infection prevalence 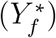 to the alternative host competition parameter (*α*_*fa*_) is

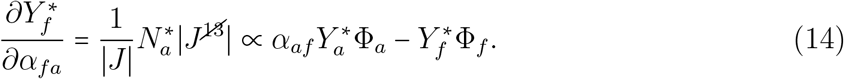

The sensitivity can take on either sign, meaning that increased competitive ability of the alternative host increases or decreases focal host infection prevalence when the focal host is a sink (Φ_*f*_ < 0) or source (Φ_*f*_ > 0), respectively. However, if the focal host is a sufficiently strong interspecific competitor (*α*_*af*_ large), then the response can be reversed when the alternative host is a sufficiently large sink (Φ_*a*_ < 0) or source (Φ_*a*_ > 0), respectively.

The global sensitivities show that greater competitive ability of the alternative can have two effects. First, greater competitive ability of the alternative host increases focal host infection prevalence at all time points when *R*_0*a*_ is sufficiently larger than *R*_0*f*_ . This effect is strongly influential ( ∣ PRCC∣ ≥ 0.5) only when interspecific competition is sufficiently strong (*α*_*af*_ *α*_*fa*_ sufficiently large within dashed maroon region in Figure 4A). Second, greater competitive ability of the alternative host decreases focal host infection prevalence at all time points when *R*_0*a*_ is similar to or lower than *R*_0*f*_ . The exact condition when this effect is strongly influential is highly context dependent (details provided in Appendix Section S5.7), but this effect is never influential when *R*_0*f*_ is much larger than *R*_0*a*_ (dotted gray region does not include bottom right box in Figure 4C).

#### 4.2.8 Summary of Question 2 Findings

For the alternative host disease parameters (*χ*_*a*_, *u*_*a*_, *β*_*a*_, *ρ*_*a*_, *µ*_*a*_) and birth rate (*b*_*a*_), our global sensitivity results show that focal host infection prevalence is typically higher at all time points when the alternative host has a higher *R*_0*a*_ (larger *χ*_*a*_ and *β*_*a*_ and smaller *u*_*a*_, *µ*_*a*_, *ρ*_*a*_, and *b*_*a*_). Biologically, this means that removal of the alternative host typically causes larger decreases (increases) in focal host infection prevalence when the alternative host has a greater (lesser) ability to transmit the pathogen. Exceptions occur when *R*_0*a*_ is much less than *R*_0*f*_, such that focal host infection prevalence decreases with decreases in the alternative host mortality rate (*µ*_*a*_) and birth rate (*b*_*a*_). For the alternative host competitive ability, we find that focal host infection prevalence decreases with higher alternative host competitive ability (*α*_*fa*_), unless *R*_0*a*_ is much greater than *R*_0*f*_, where it instead increases with higher competitive ability. These general patterns and exceptions qualitatively agree with the local sensitivity results, but the local sensitivity formulas show that exceptions can arise under additional conditions; we will address this in the Discussion.

We note two general patterns about when the parameters are strongly influential (Figure 4AC). First, the shedding (*χ*_*a*_), uptake (*u*_*a*_), and birth (*b*_*a*_) rates and interspecific competitive ability (*α*_*fa*_) of the alternative host are strongly influential across parameter space (dotted gray, long dashed purple, and long dashed orange regions). The only exception is that the shedding and uptake rates and interspecific competitive ability are not strongly influential when the alternative host’s ability to transmit the pathogen is much lower than the focal host (bottom right of Figure 4C). Second, the other disease parameters are strongly influential over much smaller regions of parameter space (short dashed blue for *β*_*a*_, solid green for *u*_*a*_, and no region for *ρ*_*a*_). For example, the transmission coefficient is only influential when competition is not too strong and the alternative host has a sufficiently high *R*_0*a*_ (long dashed blue region).

### 4.3 Question 3. Which alternative host traits have the largest influence on how quickly focal host infection prevalence changes?

Here, we use global sensitivity analysis to determine how the alternative host traits affect the divergence time. Recall that the divergence time is the amount of time needed for focal host infection prevalence to deviate from its initial value by 5% of the total change observed during alternative host loss (i.e., 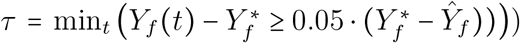). This helps identify how the traits of the alternative host influence the speed at which focal host infection prevalence changes when the alternative host begins to be lost from the disease system.

As before, we focus on identifying regions of parameter space where parameters are strongly influential ( ∣PRCC ∣ ≥0.5). Details about regions where parameters are moderately influential (0.1 ≤ ∣PRCC ∣ < 0.5) or noninfluential ( ∣ PRCC < ∣0.1) are provided in Appendix Section S6.

#### 4.3.1 Effect of Alternative Host Shedding Rate (*χ*_*a*_) on Divergence Time

Higher alternative host shedding rates (*χ*_*a*_) increase divergence time across parameter space (see Appendix Section S6.1). This effect is strongly influential ( ∣ PRCC∣ ≥ 0.5) except when *R*_0*a*_ is less than 1 and *R*_0*f*_ is large (dotted regions in Figure 4BD).

### 4.3.3 Effect of Alternative Host Uptake Rate (*u*_*a*_) on Divergence Time

Higher alternative host uptake rates (*u*_*a*_) almost always decrease divergence time. Exceptions occur when hosts do not compete against each other (*α*_*af*_ = *α*_*fa*_ = 0) and when both hosts have moderate or low *R*_0*i*_ (see Appendix Section S6.2). When hosts compete with each other, the effect is only strongly influential ( ∣ PRCC∣ ≥ 0.5) when *R*_0*a*_ is less than or similar to *R*_0*f*_ and when host competition is not too strong (dot-dashed red regions in Figure 4BD). When hosts do not compete with each other, the effect is never strongly influential (∣ PRCC∣ ≥ 0.5; dot-dashed red region does not include when *α*_*fa*_ = *α*_*af*_ = 0 in Figure 4B).

#### 4.3.3 Effect of Alternative Host Transmission Coefficient (*β*_*a*_) on Divergence Time

Higher alternative host transmission coefficients (*β*_*a*_) increase divergence time (see Appendix Section S6.3). This effect is only strongly influential (∣PRCC∣ ≥ 0.5) when *R*_0*f*_ = 2.1, *R*_0*a*_ is greater than or similar to *R*_0*f*_, and interspecific competition is not too strong (*α*_*fa*_*α*_*af*_ sufficiently small; short dashed blue regions in Figure 4BD).

#### 4.3.4 Effect of Alternative Host Recovery Rate (*ρ*_*a*_) on Divergence Time

Higher alternative host recovery rates (*ρ*_*a*_) decrease divergence time (see Appendix Section S6.4). However, this effect is never strongly influential (∣ PRCC ∣ ≥0.5; *ρ*_*a*_ has no regions in Figure 4BD).

#### 4.3.5 Effect of Alternative Host Disease Mortality Rate (*µ*_*a*_) on Divergence Time

Higher alternative host disease mortality rates (*µ*_*a*_) decrease divergence time (see Appendix Section S6.5). This effect is only strongly influential ( ∣ PRCC∣ ≥ 0.5) when the focal host is a strong competitor (*α*_*af*_ = 1.2) and *R*_0*a*_ is much higher than *R*_0*f*_ (solid green regions in Figure 4BD).

#### 4.3.6 Effect of Alternative Host Birth Rate (*b*_*a*_) on Divergence Time

Higher alternative host birth rates (*b*_*a*_) almost always increase divergence time. This effect is strongly influential across parameter space, except in idiosyncratic cases where the hosts have similar *R*_0*i*_ values (dotted gray region in Figure 4B and long dashed orange region in Figure 4D). Higher alternative host birth rates can decrease divergence time when the focal host is a weak competitor (*α*_*af*_ small), the alternative host is a moderately strong competitor (*α*_*fa*_ *=* 0.6), and both host have *R*_0*i*_ values near 2.1. However, this effect is not strongly influential.

#### 4.3.7 Effect of Alternative Host Competitive Ability (*α*_*fa*_) on Divergence Time

Higher alternative host competitive ability (*α*_*fa*_) increases divergence time when hosts have sufficiently different *R*_0*i*_ values, while it decreases divergence time when hosts have similar *R*_0*i*_ values. These effects are strongly influential in many regions of parameter space (long dashed purple in Figure 4B and dotted gray in 4D; additional details are provided in Appendix Section S5.7). First, higher alternative host competitive ability increases divergence time when the alternative host has a much higher *R*_0*i*_ than the focal host (top left block of each 3-by-2 grid); this effect is strongly influential except when both hosts are sufficiently weak competitors (*α*_*fa*_ and *α*_*af*_ sufficiently small). Second, higher alternative host competitive ability also increases divergence time when the alternative host has a much lower *R*_0*i*_ than the focal host (bottom-right three blocks of each 3-by-2 grid); this effect is strongly influential only when the alternative host is a sufficiently strong interspecific competitor (*α*_*fa*_ sufficiently large). Third, higher alternative host competitive ability decreases divergence time when the alternative and focal hosts have similar *R*_0*i*_ values (middle left and top right blocks of each 3-by-2 grid); this effect is strongly influential only when the focal host is a sufficiently weak interspecific competitor (*α*_*af*_ sufficiently small). Altogether, the negative and positive effects of alternative host competitive ability are strongly influential for many combinations of host *R*_0*i*_ values and interspecific competition coefficients, but no positive or negative effect is strongly influential for all combinations of the *R*_0*i*_ and *α*_*ij*_ values.

#### 4.3.8 Summary of Question 3 Findings

Across parameter space, we find that higher *R*_0*a*_ (higher *χ*_*a*_, *β*_*a*_, and lower *u*_*a*_, *µ*_*a*_, *ρ*_*a*_), higher birth rates (*b*_*a*_), and higher competitive ability (*α*_*fa*_) typically increase the divergence time. In other words, focal host infection prevalence responds faster to the loss of the alternative host when the alternative host has a greater ability to transmit disease, has higher reproduction rates, and is a stronger interspecific competitor. Exceptions occur when (i) hosts do not compete and both host have low *R*_0*i*_ values, reversing the effect of *u*_*a*_, and (ii) when hosts have similar *R*_0*i*_ values, reversing the effect of *α*_*fa*_.

We note two general patterns about when the parameters are strongly influential (Figure 4BD).

First, the shedding (*χ*_*a*_) and birth (*b*_*a*_) rates and interspecific competitive ability (*α*_*fa*_) of the alternative host are strongly influential across parameter space (dotted gray and long dashed purple regions). One caveat is that shedding rates and competitive ability are not strongly influential when the alternative host’s ability to transmit the pathogen is much lower than the focal host (e.g., *R*_0*a*_ = 0.7 and *R*_0*f*_ = 6.3; bottom right of Figure 4C). A second caveat is that, as explained in the previous subsection, the effect of interspecific competitive ability depends on the exact values of the *R*_0*i*_ and competition coefficients. Second, the other disease parameters are strongly influential over much smaller regions of parameter space (dot-dashed red for *u*_*a*_, short dashed blue for *β*_*a*_, solid green for *µ*_*a*_, and no region for *rho*_*a*_). For example, the transmission coefficient is only influential when interspecific competition is sufficiently weak (short dashed blue in Figure 4B) and the alternative host has a sufficiently high *R*_0*a*_ (short dashed blue in Figure 4D).

## 5 Discussion

In this study, we used local and global sensitivity analysis to explore the impacts of alternative host trait values on the changes in focal host infection prevalence as the alternative host is lost from the system. First, we assessed how alternative host traits influence the changes in focal host infection prevalence. We found three types of responses: (i) monotonic decreases when *R*_0*a*_ is sufficiently high relative to *R*_0*f*_ (Figure 3F), (ii) monotonic increases when *R*_0*a*_ is sufficiently low relative to *R*_0*f*_ (Figure 3F), and (iii) nonmonotonic changes when the hosts had similar *R*_0*i*_ values and alternative host competitive ability was sufficiently high (*α*_*fa*_ large; Figure 3F). Second, we determined which traits have the largest influence on whether focal host infection prevalence is higher or lower during transient dynamics. We found that focal host infection prevalence is typically larger when the alternative host has a higher *R*_0*i*_ value (larger *χ*_*a*_ and *β*_*a*_, smaller *u*_*a*_, *µ*_*a*_, *ρ*_*a*_), lower birth rate (*b*_*a*_), or lower competitive ability (*α*_*fa*_). However, exceptions occur for *µ*_*a*_ and *b*_*a*_ when *R*_0*a*_ is much less than *R*_0*f*_ and for *α*_*fa*_ when *R*_0*a*_ is much greater than *R*_0*f*_ . Additionally, we found that shedding (*χ*_*a*_), uptake (*u*_*a*_), birth (*b*_*a*_) rates and interspecific competitive ability (*α*_*fa*_) of the alternative host are strongly influential across parameter space (Figure 4AC). In contrast, alternative host disease mortality rate *µ*_*a*_ and recovery rate *ρ*_*a*_ were rarely influential, and the alternative host transmission coefficient *β*_*a*_ was only influential when *R*_0*a*_ was similar or greater than *R*_0*f*_ (Figure 4AC). Finally, we assessed the influence of alternative host traits on how quickly focal host infection prevalence diverges from its initial value. We found that higher *R*_0*a*_, birth rate, and competitive ability all increase the divergence time, with exceptions occurring when *R*_0*a*_ is similar to *R*_0*f*_ and when alternative hosts do not compete. Similar to the above, we found that alternative host shedding rates (*χ*_*a*_), birth rate (*χ*_*a*_), and competitive ability (*α*_*fa*_) were strongly influential throughout most of parameter space whereas the other parameters were strongly influential in much smaller regions of parameter space (Figure 4BD). Here, we explain how results can be used by empiricists and how our results connect to the literature on amplification and dilution effects.

Our theoretical results offer two general predictions. First, when predicting how focal host infection prevalence will respond to alternative host loss, our results show that measuring each host’s ability to transmit the pathogen (*R*_0*i*_) is typically more important than measuring their competitive abilities. This is because the loss of the alternative host monotonically increases and monotonically decreases focal host infection prevalence when *R*_0*a*_ is much less than and much greater than *R*_0*f*_, respectively (orange cells where *R*_0*a*_<< *R*_0*f*_ and blue cells where *R*_0*a*_ >> *R*_0*f*_ in Figure 3F). It is only when the hosts have similar abilities to transmit the disease (*R*_0*f*_ similar to *R*_0*a*_) that measuring host competitive ability becomes important for determining if focal host infection prevalence will decrease monotonically, increase monotonically, or change nonmonotonically (cells change from blue to orange with increasing *α*_*fa*_ and red dots occur only when *R*_0*a*_ ≈ *R*_0*f*_ and *α*_*fa*_ is sufficiently large in Figure 3F). Second, when estimating an alternative host’s ability to transmit the pathogen, our results show that estimating host shedding rates (*χ*_*i*_), uptake rates (*u*_*i*_), and birth rate (*b*_*a*_) is generally more important than estimating other parameters, including the transmission parameter (*β*_*a*_), disease mortality rate (*µ*_*a*_), and recovery rate (*ρ*_*a*_). In total, while accurate estimation of all parameters can be necessary, our results identify which traits should be prioritized to make qualitatively and quantitatively accurate predictions about how focal host infection prevalence will respond to alternative host loss.

The effects of host losses on focal host disease levels are typically discussed in terms of dilution (disease levels increase with host loss) or amplification (disease decrease with host loss) [Keesing et al., 2006, Wood and Lafferty, 2013]. In this context, our results show that the alternative host consistently amplifies and dilutes disease in cases where its loss causes focal host infection prevalence to monotonically increase or decrease, respectively (blue and orange in Figure 3F). In contrast, the effects of the alternative host do not cleanly fall into one category (i.e., amplification or dilution) when its loss causes nonmonotonic changes in focal host infection prevalence (red dots in Figure 3F). Keesing et al. [2006] identified five mechanisms through which an alternative host affects focal host disease levels. Translating those mechanisms to environmentally transmitted pathogens [Johnson and Thieltges, 2010], the patterns we observe are driven by the combined effects of two mechanisms: host-parasite encounter reduction and susceptible host regulation. The first mechanism (host-parasite encounter reduction) appears in our model because higher *R*_0*i*_ values lead to increases in spore densities, which increases host-parasite contact rates. The second mechanism (susceptible host regulation) appears in our model because increases in alternative host competitive abilities (*α*_*fa*_) reduce focal host density. When interpreted in terms of these two mechanisms, our results suggest that host-parasite encounter reduction is often more important than susceptible host regulation when predicting the responses of focal host infection prevalence to loss of the alternative. However, susceptible host regulation becomes more important when the host species have similar abilities to transmit the pathogen.

We have quantified a host’s ability to spread disease using the pathogen’s basic reproductive number (*R*_0*i*_) and the net effect of a host species on propagule density (Φ_*i*_). However, prior studies have argued that a more appropriate metric is host competence, i.e., the ability of an exposed individual to transmit the pathogen [Stewart Merrill et al., 2022, Mommer et al., 2026]. Stewart Merrill and Johnson [2020] have argued that competence should be a host trait that is distinct from the effect of host densities on pathogen exposure, instead comprising of only infection, parasite development, and pre-transmission survival.

For our model, the competence of host *i* is

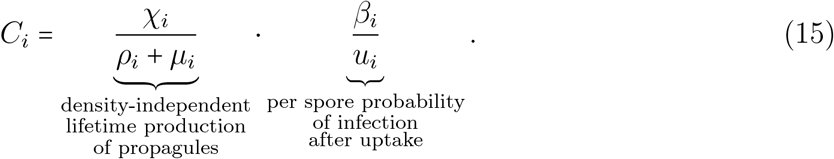

This is biologically interpreted as the product of the per-spore probability of infection after uptake and the lifetime production of propagules by infected individuals, assuming density-independent mortality. Combined, this product is the total number of propagules produced by an exposed individual before recovery or death, in the absence of density-dependent mortality. While the relative orderings of the *R*_0*i*_, Φ_*i*_, and *C*_*i*_ values do not have to agree [Cortez, 2024], host species were always ordered in the same way by all three metrics of host disease-spreading capacity across all nominal parameter sets. That is, the host species with a larger *R*_0*i*_ value also had a larger Φ_*i*_ and *C*_*i*_ value. This means that all of our results about *R*_0*a*_ being larger than, equal to, or smaller than *R*_0*f*_ can also be interpreted in terms of the alternative host has higher, equal, or lower competence than the focal host, respectively.

We used both local and global sensitivity analysis so that we could draw on the strengths of both approaches. While the analytical local sensitivity analysis formulas only necessarily apply at the two-host endemic equilibrium, the main advantage of that approach is that the formulas apply across parameter space. Our local sensitivity results agree with those of previous work [Searle et al., 2016, Cortez, 2021, Cortez and Duffy, 2021, Cortez, 2024]. Specifically, alternative host shedding rate (*χ*_*a*_) and uptake rate (*u*_*a*_) have a positive and negative effect, respectively, across all of parameter space, while the transmission coefficient (*β*_*a*_) and mortality rate (*µ*_*a*_) have a positive and negative effect, respectively, in much of parameter space, but can switch signs depending on the competitive strength and competence of each host [Cortez, 2021, Cortez and Duffy, 2021, Cortez, 2024]. Overall, this supports the general prediction from prior equilibrium-based studies that higher and lower competence alternative hosts typically increase and decrease, respectively, disease risk and disease levels in a focal host, but those effects can be reversed if the alternative host is a sufficiently strong competitor [Johnson et al., 2013, Mihaljevic et al., 2014, O’Regan et al., 2015, Rudolf and Antonovics, 2005, Searle et al., 2016, Cortez, 2024, Dobson, 2004, Fenton et al., 2015, Espira et al., 2022].

Global sensitivity analysis, on the other hand, was numerically calculated and thus provided no analytical formulas. However, the global sensitivities quantified how the influence of each alternative host parameter varied through time and determined the relative strengths of the effects of each alternative host parameter. We found that our local and global sensitivity results often agreed, but note two exceptions. First, for most nominal parameter sets, the local sensitivities at the two-host endemic equilibrium had the same signs as the global sensitivities at all time points. The only exception was the alternative host birth rate (*b*_*a*_), where an increase in *b*_*a*_ typically decreased focal host infection prevalence at the two-host endemic equilibrium, but it decreased focal host infection prevalence during the transient dynamics for one nominal parameter set. Although the parameter was not strongly influential ( ∣ PRCC∣ < 0.5) in that region of parameter space, this highlights an important limitation of current equilibrium-based dilution-effect theory to explain effect on transient disease dynamics. Second, some of the effects of parameters predicted by the local sensitivity formulas were not observed in our global sensitivity results. Specifically, the local sensitivity formulas for the alternative host transmission parameter (*β*_*a*_) and recovery rate (*ρ*_*a*_) predict that both parameters can have positive or negative effects on focal host infection prevalence at the two-host endemic equilibrium. In contrast, our numerical global sensitivity results only showed that the transmission parameter had a positive effect and the recovery rate had a negative effect. These differences suggest that some of our predictions may not hold in extreme regions of parameter space where differences in host competence or competitive ability between the two species are very large. Altogether, this study illustrates how combining local and global sensitivity approaches can be provide a more holistic understanding of the effects of an alternative host on focal host infection prevalence.

While our study provides general rules about how alternative host traits influence the dynamics of focal host infection prevalence, our results are limited by specific assumptions built into our model. In model (1), we made simplifying assumptions that are relaxed in other environmental transmission models, including density-independent reproduction rates [Chen et al., 2022, Searle et al., 2016, Cortez, 2021], infection rates with a bilinear form [Joh et al., 2009, Cortez and Weitz, 2013, Breban, 2013, Clay et al., 2021], and birth rates, uptake rates, and competitive abilities that are independent of infection status [Searle et al., 2016, Cáceres et al., 2014]. We made these assumptions to reduce the number of parameters in our system, but relaxing those assumptions could yield additional insight into how host traits shape disease dynamics. Our approach also leaves room for future work to investigate two-host communities composed of hosts with mutualistic or predator-prey interactions [Su et al., 2022, Hatcher et al., 2006, Lopez et al., 2023]. Finally, while we investigated the transient dynamics that occur during host loss, understanding how host traits affect disease dynamics after host invasions or additions is also an important area of future work [Hanthanan Arachchilage et al., 2023].

In summary, this study used local and global sensitivity analysis to explore how the transient disease dynamics in a focal host are affected by the loss of an alternative host from the system. Our results provide predictions about which and how traits of the alternative host strongly influence the dynamics of the alternative host. This study broadens our understanding of how host species can impact each other’s disease levels.

## 6 Acknowledgments

MHC was supported by the National Science Foundation under Award DEB-2015280. MLR thanks FSU for support.

