## Supplemental Appendices S1-S6 for "Host traits predict the transient disease dynamics that occur during the loss of an alternative host"

### Supplement

Miles L. Roberston & Michael H. Cortez

October 29, 2026

**S3 Numerical Simulation of Nominal Parameter Sets** **13**

**S4 Key for Global and Local Sensitivity Analysis Results** **15**

**S5 Focal Host Infection Prevalence: Simulation and Global & Local Sensitivity Analysis Results** **15**

S6.7 Alternative Host Competitive Ability ($\alpha_{fa}$) Effect on Divergence Time 37

### S1 Model Proofs

Our nondimensionalized $NYP$ model is:

$$\begin{aligned}
\frac{\mathrm{d}N_f}{\mathrm{d}t} &= \underbrace{b_f N_f}_{\text{reproduction and mortality}} - \underbrace{(N_f + \alpha_{fa} N_a) N_f}_{\substack{\text{non-disease mortality} \\ \text{due to intra- and} \\ \text{interspecific interactions}}} - \underbrace{\mu_f Y_f N_f}_{\text{disease mortality}} \\
\frac{\mathrm{d}N_a}{\mathrm{d}t} &= \underbrace{b_a N_a}_{\text{reproduction and mortality}} - \underbrace{(N_a + \alpha_{af} N_f) N_a}_{\substack{\text{non-disease mortality} \\ \text{due to intra- and} \\ \text{interspecific interactions}}} - \underbrace{\mu_a Y_a N_a}_{\text{disease mortality}} \\
\frac{\mathrm{d}Y_f}{\mathrm{d}t} &= \underbrace{(1 - Y_f) P}_{\text{transmission}} - \underbrace{b_f Y_f}_{\text{reproduction}} - \underbrace{\rho_f Y_f}_{\text{recovery}} - \underbrace{\mu_f Y_f (1 - Y_f)}_{\text{disease mortality}} \\
\frac{\mathrm{d}Y_a}{\mathrm{d}t} &= \underbrace{\beta_a (1 - Y_a) P}_{\text{transmission}} - \underbrace{b_a Y_a}_{\text{reproduction}} - \underbrace{\rho_a Y_a}_{\text{recovery}} - \underbrace{\mu_a Y_a (1 - Y_a)}_{\text{disease mortality}} \\
\frac{\mathrm{d}P}{\mathrm{d}t} &= \underbrace{\chi_f Y_f N_f + \chi_a Y_a N_a}_{\text{spore release}} - \underbrace{(u_f N_f + u_a N_a) P}_{\text{uptake}} - \underbrace{P}_{\text{degradation}} .
\end{aligned} \tag{S1}$$

#### S1.1 Focal Host-Only Endemic Equilibrium

The $NYP$ form of the focal host-only system is defined by

$$\begin{aligned}
\frac{\mathrm{d}N_f}{\mathrm{d}t} &= b_f N_f - N_f^2 - \mu_f Y_f N_f \\
\frac{\mathrm{d}Y_f}{\mathrm{d}t} &= (1 - Y_f) P - (b_f + \rho_f) Y_f - \mu_f Y_f (1 - Y_f) \\
\frac{\mathrm{d}P}{\mathrm{d}t} &= \chi_f Y_f N_f - u_f N_f P - P.
\end{aligned} \tag{S2}$$

We use the $NIP$ version of this model to prove the unique existence of the endemic equilibrium, which is

$$\begin{aligned}
\frac{\mathrm{d}N_f}{\mathrm{d}t} &= b_f N_f - N_f^2 - \mu_f I_f \\
\frac{\mathrm{d}I_f}{\mathrm{d}t} &= S_f P - N_f I_f - (\rho_f + \mu_f) I_f
\end{aligned}$$

$$\frac{\mathrm{d}P}{\mathrm{d}t} = \chi_f I_f - u_f N_f P - P.$$

**Lemma SA1:** A unique endemic equilibrium exists if and only if $R_{0f} > 1$ and $\chi_f > \mu_f u_f$.

From Maple, we find the following endemic equilibrium value for the focal host-only $NIP$ model:

$$\text{Intermediate Values: } \begin{cases} K = \rho_f + \mu_f \\ E = \chi_f - u_f \mu_f \\ F = b_f - \mu_f \\ \Omega = \mu_f(K u_f + 1) + \chi_f F \\ \Delta = (\Omega - 2\mu_f)^2 + 4\mu_f \chi_f (K + F) \\ \Theta = \chi_f - u_f(b_f + K) - 1 \end{cases}$$

$$N_f^* = (\Omega + \sqrt{\Delta})/(2E)$$

$$I_f^* = (\Theta N_f^* - K)/E$$

$$P^* = ((\chi_f - b_f u_f - 1)N_f^* - K(b_f u_f + 1))/(1 + u_f(b_f - \mu_f))$$

From $\frac{\mathrm{d}P}{\mathrm{d}t} = 0$, if $N_f^*$ and $I_f^*$ are positive, $P^*$ is also positive.

$N_f^*$ is the larger of two real roots of a quadratic equation, which are always real as $\Delta$ is always positive. These two roots are both negative if $\chi_f < u_f \mu_f$, and exactly one is positive (the one written above as $N_f^*$) if $\chi_f > u_f \mu_f$. Therefore, we assume $\chi_f > u_f \mu_f$ (and therefore $E > 0$) for all focal host-only endemic equilibria.

Since $E > 0$, $I_f^*$ is positive if its numerator is. This occurs if $\sqrt{\Delta} > (2EK)/\Theta - \Omega$. If $\sqrt{\Delta} > |(2EK)/\Theta - \Omega|$, then $I_f^*$ is positive. This inequality implies and is implied by $\Theta b_f - K > 0$, which also implies and is implied by $R_{0f} > 1$ at the focal host disease-free equilibrium: $\hat{N}_f = b_f$. Additionally, parameter sets do not exist where $-\sqrt{\Delta} > (2EK)/\Theta - \Omega$, as this implies $K < \Theta((\Omega - \sqrt{\Delta})/(2E))$, the parenthetical term being the always-negative solution to the quadratic equation that gave $N_f^*$.

Therefore, a single endemic equilibrium exists if $\chi_f > u_f\mu_f$ and $R_{0f}(\hat{N}_f) > 1$, and none exist otherwise. □

To investigate the stability of this endemic equilibrium, we randomly generated 10 million parameter sets (all parameters sampled from $U(0, 10)$, except $\chi_f$, which was sampled from $U(0, 100)$) and discarded those that did not satisfy $R_{0f} > 1$ and $\chi_f > u_f\mu_f$, leaving over 3 million remaining. Some of those parameter sets did give an unstable endemic equilibrium (e.g., $(b_f, \rho_f, \mu_f, \chi_f, u_f) = (0.76966, 0.02664, 3.32406, 90.48979, 1.42789)$, which gave Jacobian eigenvalues $-4.92319$, $0.00106 - 0.57609i$, and $0.00106 + 0.57609i$). However, focal host endemic equilibrium instability occurred in less than 0.006% of parameter sets satisfying $R_{0f} > 1$ and $\chi_f > u_f\mu_f$, and only occurred when $\chi_f$ was 30 times greater than all other parameters and $\mu_f \gg b_f$, neither of which occurred in or near any nominal parameter sets.

#### S1.2 Two-Host Disease-Free Equilibrium

When the disease is absent from the system, the model simplifies to

$$\frac{\mathrm{d}N_f}{\mathrm{d}t} = b_f N_f - (N_f + \alpha_{fa} N_a) N_f$$
$$\frac{\mathrm{d}N_a}{\mathrm{d}t} = b_a N_a - (N_a + \alpha_{af} N_f) N_a.$$

The equilibrium where both hosts coexist occurs at

$$\tilde{N}_f = (b_f - \alpha_{fa} b_a)/(1 - \alpha_{fa}\alpha_{af})$$
$$\tilde{N}_a = (b_a - \alpha_{af} b_f)/(1 - \alpha_{af}\alpha_{fa}).$$

From Maple, we find that $\tilde{N}_f$ and $\tilde{N}_a$ are both positive and form a stable equilibrium if and only if

$$b_f > \alpha_{fa} b_a$$
$$b_a > \alpha_{af} b_f$$
$$1 > \alpha_{af}\alpha_{fa}$$

are all true. Therefore, if $1 < \alpha_{af}\alpha_{fa}$, the two-host coexistence disease-free equilibrium is not stable.

#### S1.3 Full System Jacobian

The Jacobian for (S1) is:

$$J = \begin{bmatrix} \begin{matrix} b_f \\ -(2N_f + \alpha_{fa} N_a) \\ -\mu_f Y_f \end{matrix} & -\alpha_{fa} N_f & -\mu_f N_f & 0 & 0 \\ -\alpha_{af} N_a & \begin{matrix} b_a \\ -(2N_a + \alpha_{af} N_f) \\ -\mu_a Y_a \end{matrix} & 0 & -\mu_a N_a & 0 \\ 0 & 0 & \begin{matrix} -P - (b_f + \rho_f) \\ -\mu_f (1 - Y_f) \\ +\mu_f Y_f \end{matrix} & 0 & 1 - Y_f \\ 0 & 0 & 0 & \begin{matrix} -\beta_a P - (b_a + \rho_a) \\ -\mu_a (1 - Y_a) + \mu_a Y_a \end{matrix} & \beta_a (1 - Y_a) \\ \chi_f Y_f - u_f P & \chi_a Y_a - u_a P & N_f \chi_f & N_a \chi_a & -(u_f N_f + u_a N_a + 1) \end{bmatrix} \tag{S3}$$

Note that, at the two-host endemic equilibrium, the $\partial \dot{P} / \partial N_i$ entry for each host can be written in terms of the source/sink metric ($\Phi_i = \chi_i - u_i P^* / Y_i^*$) used in local sensitivity analysis: $\partial \dot{P} / \partial N_i = Y_i^* \Phi_i$.

#### S1.4 $R_{0f}$ and $R_{0a}$ Calculation

For each host, we calculate the $R_{0i}$ value with the next generation method: we calculate the Jacobian for the one-host disease model, filter out rows and columns not associated with a disease state variable (remove $N_i$, retain $Y_i$ and $P$), assemble the transmission matrix ($T$) and the transition matrix ($F$), calculate the spectral radius of $-TF^{-1}$, and evaluate it at the one-host disease-free equilibrium $\hat{N}_i = b_i$. Specifically,

$$T = \begin{bmatrix} 0 & 0 \\ \hat{N}_i \chi_i & 0 \end{bmatrix} \quad F = \begin{bmatrix} -b_i - \rho_i - \mu_i & \beta_i \\ 0 & -\hat{N}_i u_i - 1 \end{bmatrix}$$

$$R_{0i}(\hat{N}_i) = \frac{\beta_i \chi_i \hat{N}_i}{(b_i + \rho_i + \mu_i)(u_i \hat{N}_i + 1)} \implies R_{0i}(b_i) = \frac{\beta_i \chi_i b_i}{(b_i + \rho_i + \mu_i)(u_i b_i + 1)}.$$

This gives

$$R_{0a}(b_a) = \frac{\beta_a \chi_a b_a}{(b_a + \rho_a + \mu_a)(u_a b_a + 1)} \quad \text{and} \quad R_{0f}(b_f) = \frac{\chi_f b_f}{(b_f + \rho_f + \mu_f)(u_f b_f + 1)}.$$

### S2 Calculations of Local Sensitivity Analysis

#### S2.1 Two-Host Endemic Equilibrium Model

Here, we assume that $\zeta_a$ is small enough that $\zeta_a t$ is effectively constant with respect to all other system dynamics, replacing it with $\eta_a$.

$$\begin{aligned}
\frac{\mathrm{d}N_f}{\mathrm{d}t} &= b_f N_f - (N_f + \alpha_{fa} N_a) N_f - \mu_f I_f \\
\frac{\mathrm{d}N_a}{\mathrm{d}t} &= (b_a - \eta_a) N_a - (N_a + \alpha_{af} N_f) N_a - \mu_a I_a \\
\frac{\mathrm{d}I_f}{\mathrm{d}t} &= S_f P - (N_f + \alpha_{fa} N_a) I_f - (\rho_f + \mu_f) I_f \\
\frac{\mathrm{d}I_a}{\mathrm{d}t} &= \beta_a S_a P - (N_a + \alpha_{af} N_f) I_a - (\rho_a + \mu_a + \eta_a) I_a \\
\frac{\mathrm{d}P}{\mathrm{d}t} &= \chi_f I_f + \chi_a I_a - (u_f N_f + u_a N_a) P - P
\end{aligned}$$

We convert this model to tracking infection prevalence by using $Y_f = I_f / N_f$ and $Y_a = I_a / N_a$:

$$\begin{aligned}
\frac{\mathrm{d}N_f}{\mathrm{d}t} &= b_f N_f - (N_f + \alpha_{fa} N_a) N_f - \mu_f Y_f N_f \\
\frac{\mathrm{d}N_a}{\mathrm{d}t} &= (b_a - \eta_a) N_a - (N_a + \alpha_{af} N_f) N_a - \mu_a Y_a N_a \\
\frac{\mathrm{d}Y_f}{\mathrm{d}t} &= (1 - Y_f) P - (N_f + \alpha_{fa} N_a) Y_f - (\rho_f + \mu_f) Y_f - \frac{Y_f}{N_f} \frac{\mathrm{d}N_f}{\mathrm{d}t}
\end{aligned}$$

$$\frac{\mathrm{d}Y_a}{\mathrm{d}t} = \beta_a(1-Y_a)P - (N_a + \alpha_{af}N_f)Y_a - (\rho_a + \mu_a + \eta_a)Y_a - \frac{Y_a}{N_a}\frac{\mathrm{d}N_a}{\mathrm{d}t}$$
$$\frac{\mathrm{d}P}{\mathrm{d}t} = \chi_f Y_f N_f + \chi_a Y_a N_a - (u_f N_f + u_a N_a)P - P$$

In many cases, avoiding expansion of the derivatives in the $Y$ equations is wise. However, in this case, expanding this term removes dependence of $Y$ on the $N$ state variables:

$$\frac{\mathrm{d}N_f}{\mathrm{d}t} = b_f N_f - (N_f + \alpha_{fa}N_a)N_f - \mu_f Y_f N_f$$
$$\frac{\mathrm{d}N_a}{\mathrm{d}t} = (b_a - \eta_a)N_a - (N_a + \alpha_{af}N_f)N_a - \mu_a Y_a N_a$$
$$\frac{\mathrm{d}Y_f}{\mathrm{d}t} = (1-Y_f)P - (\rho_f + b_f)Y_f - \mu_f Y_f(1-Y_f)$$
$$\frac{\mathrm{d}Y_a}{\mathrm{d}t} = \beta_a(1-Y_a)P - (\rho_a + b_a)Y_a - \mu_a Y_a(1-Y_a)$$
$$\frac{\mathrm{d}P}{\mathrm{d}t} = \chi_f Y_f N_f + \chi_a Y_a N_a - (u_f N_f + u_a N_a)P - P$$

At any equilibrium, the following system of equations is satisfied. Note that the $N$ equations have a factor of total host density divided from it, and the $Y$ equations have a factor of $(1 - Y)$ divided from it (the latter ensures more entries have a fixed sign). This system of equations will be used to calculate the sensitivity of $Y_f^*$ with respect to secondary host parameters.

$$\begin{aligned}
0 &= b_f - (N_f^* + \alpha_{fa}N_a^*) - \mu_f Y_f^* \\
0 &= (b_a - \eta_a) - (N_a^* + \alpha_{af}N_f^*) - \mu_a Y_a^* \\
0 &= P^* - (\rho_f + b_f)Y_f^*/(1-Y_f^*) - \mu_f Y_f^* \\
0 &= \beta_a P^* - (\rho_a + b_a)Y_a^*/(1-Y_a^*) - \mu_a Y_a^* \\
0 &= \chi_f Y_f^* N_f^* + \chi_a Y_a^* N_a^* - (u_f N_f^* + u_a N_a^*)P^* - P^* \qquad \text{(S4)}
\end{aligned}$$

#### S2.2 Jacobian of Model

The Jacobian for (S4) is

$$J = \begin{bmatrix} -1 & -\alpha_{fa} & -\mu_f & 0 & 0 \\ -\alpha_{af} & -1 & 0 & -\mu_a & 0 \\ 0 & 0 & J_{33} & 0 & 1 \\ 0 & 0 & 0 & J_{44} & \beta_a \\ Y_f^* \Phi_f & Y_a^* \Phi_a & \chi_f N_f^* & \chi_a N_a^* & -u_f N_f^* - u_a N_a^* - 1 \end{bmatrix}$$

where $J_{33} = -\mu_f - (b_f + \rho_f)/(1 - Y_f^*)^2$, $J_{44} = -\mu_a - (b_a + \rho_a)/(1 - Y_a^*)^2$, and $\Phi_i = \chi_i - u_i Y_i^*/P^*$.

#### S2.3 Calculation of Local Sensitivity

In accordance with Yodzis [1988], we calculate the sensitivity of $X_i^*$ to parameter $p$ with system $\frac{\mathrm{d}\vec{X},t}{\mathrm{d}=}\vec{f}(\vec{X}^*, p)$ (S4) as $\frac{\partial X_i^*}{\partial p} = -\frac{1}{\det(J)} \sum_{j=1}^{N} (-1)^{i+j} \frac{\partial f_j}{\partial p} \det(J^{\cancel{ji}})$, where $J_{3,\cdot}^{-1}$ is the third row of the inverse of the Jacobian, $det(J^c ancelji)$ is the $(j, i)$ minor of the Jacobian, and $\det(J) < 0$. Given the order of the variables as presented above, the index of $Y_f$ is $i = 3$. With a 5-dimensional system, stable equilibria will have $\det(J) < 0$. The goal of the calculations here are to determine regions of parameter space where the signs of local sensitivities are predictable.

The relevant minors are:

$$\det(J^{\cancel{13}}) = -(\mu_a + \frac{b_a + \rho_a}{(1 - Y_a^*)^2})(\alpha_{af} Y_a^* \Phi_a - Y_f^* \Phi_f)$$

$$\det(J^{\cancel{23}}) = (\mu_a + \frac{b_a + \rho_a}{(1 - Y_a^*)^2})(\alpha_{fa} Y_f^* \Phi_f - Y_a^* \Phi_a)$$

$$\det(J^{\cancel{43}}) = -(1 - \alpha_{fa}\alpha_{af})\chi_a N_a^* - \mu_a(\alpha_{fa} Y_f^* \Phi_f - Y_a^* \Phi_a)$$

$$\det(J^{\cancel{53}}) = (\mu_a + \frac{b_a + \rho_a}{(1 - Y_a^*)^2})(1 - \alpha_{fa}\alpha_{af})$$

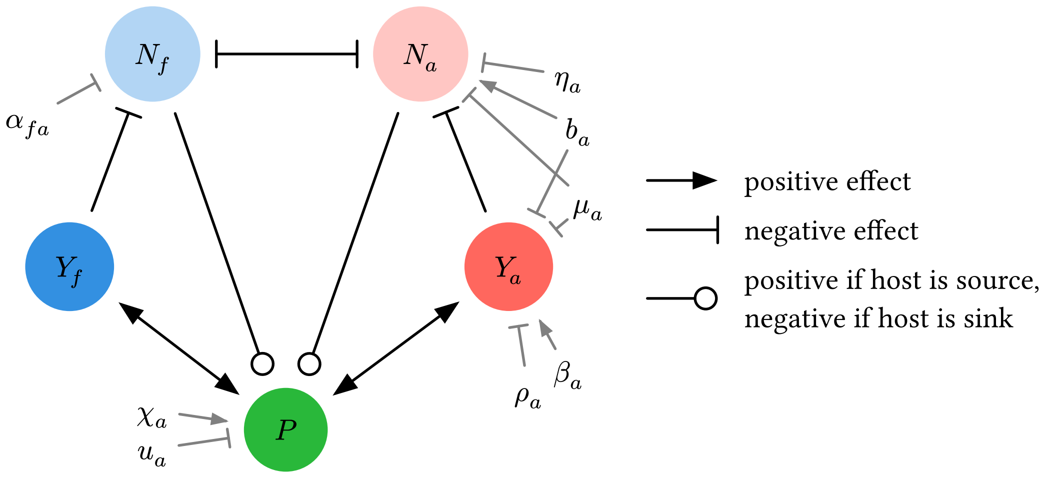

| Parameter | Pathway | Positive When | Negative When |
|---|---|---|---|
| $\chi_a$ | $\chi_a \to P \to Y_f$ | Always | Never |
| $u_a$ | $u_a \dashv P \to Y_f$ | Never | Always |
| $\beta_a$ | $\beta_a \to Y_a \to P \to Y_f$ | Always | Never |
| | $\beta_a \to Y_a \dashv N_a \dashv N_f \multimap P \to Y_f$ | $\Phi_f > 0$ | $\Phi_f < 0$ |
| | $\beta_a \to Y_a \dashv N_a \multimap P \to Y_f$ | $\Phi_a < 0$ | $\Phi_a > 0$ |
| $\rho_a$ | $\rho_a \dashv Y_a \to P \to Y_f$ | Never | Always |
| | $\rho_a \dashv Y_a \dashv N_a \dashv N_f \multimap P \to Y_f$ | $\Phi_f < 0$ | $\Phi_f > 0$ |
| | $\rho_a \dashv Y_a \dashv N_a \multimap P \to Y_f$ | $\Phi_a > 0$ | $\Phi_a < 0$ |
| $\mu_a$ | $\mu_a \dashv Y_a \to P \to Y_f$ | Never | Always |
| | $\mu_a \dashv N_a \dashv N_f \multimap P \to Y_f$ | $\Phi_f > 0$ | $\Phi_f < 0$ |
| | $\mu_a \dashv Y_a \dashv N_a \dashv N_f \multimap P \to Y_f$ | $\Phi_f < 0$ | $\Phi_f > 0$ |
| | $\mu_a \dashv N_a \multimap P \to Y_f$ | $\Phi_a < 0$ | $\Phi_a > 0$ |
| | $\mu_a \dashv Y_a \dashv N_a \multimap P \to Y_f$ | $\Phi_a > 0$ | $\Phi_a < 0$ |
| $b_a$ | $b_a \to Y_a \to P \to Y_f$ | Always | Never |
| | $b_a \to N_a \dashv N_f \multimap P \to Y_f$ | $\Phi_f < 0$ | $\Phi_f > 0$ |
| | $b_a \to Y_a \dashv N_a \dashv N_f \multimap P \to Y_f$ | $\Phi_f > 0$ | $\Phi_f < 0$ |
| | $b_a \to N_a \multimap P \to Y_f$ | $\Phi_a > 0$ | $\Phi_a < 0$ |
| | $b_a \to Y_a \dashv N_a \multimap P \to Y_f$ | $\Phi_a < 0$ | $\Phi_a > 0$ |
| $\alpha_{fa}$ | $\alpha_{fa} \dashv N_f \multimap P \to Y_f$ | $\Phi_f < 0$ | $\Phi_f > 0$ |
| | $\alpha_{fa} \dashv N_f \dashv N_a \multimap P \to Y_f$ | $\Phi_a > 0$ | $\Phi_a < 0$ |
| $\eta_a$ | $\eta_a \dashv N_a \dashv N_f \multimap P \to Y_f$ | $\Phi_f > 0$ | $\Phi_f < 0$ |
| | $\eta_a \dashv N_a \multimap P \to Y_f$ | $\Phi_a < 0$ | $\Phi_a > 0$ |

Figure S2.2: Pathway diagram demonstrating the possible effects of each alternative host parameter, given that the two hosts can coexist in the absence of the pathogen ($\alpha_{fa}\alpha_{af} < 1$). Black arrows indicate the effect of increasing one state variable on another state variable, and gray arrows indicate the effect of increasing one parameter on a state variable.

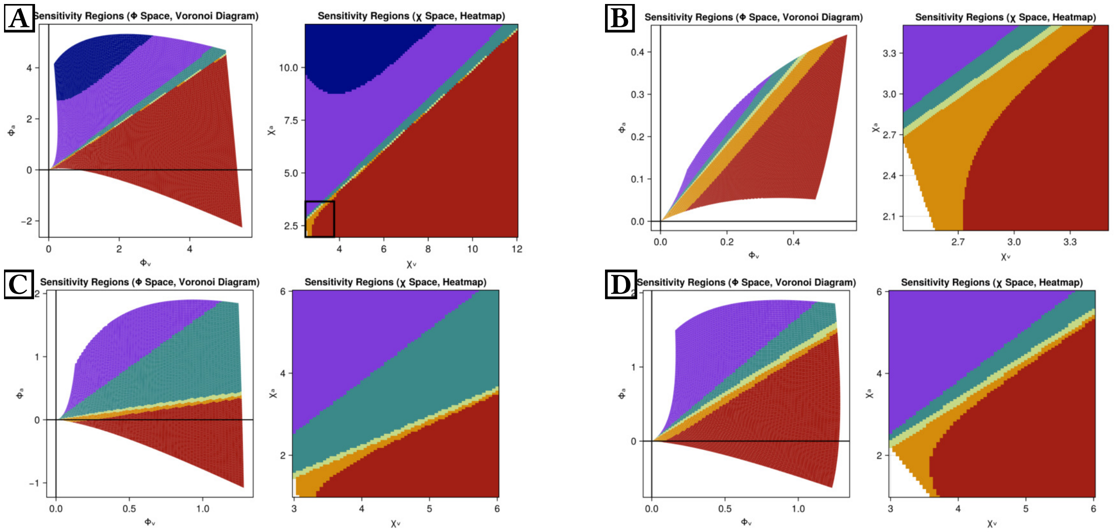

| | $\frac{\partial Y_f^*}{\partial \chi_a}$ | $\frac{\partial Y_f^*}{\partial u_a}$ | $\frac{\partial Y_f^*}{\partial \beta_a}$ | $\frac{\partial Y_f^*}{\partial \rho_a}$ | $\frac{\partial Y_f^*}{\partial \mu_a}$ | $\frac{\partial Y_f^*}{\partial b_a}$ | $\frac{\partial Y_f^*}{\partial \alpha_{fa}}$ | $\frac{\partial Y_f^*}{\partial \eta_a}$ |
|---|---|---|---|---|---|---|---|---|
| ■ (dark blue) | + | − | − | + | − | + | + | − |
| ■ (purple) | + | − | + | − | − | + | + | − |
| ■ (teal) | + | − | + | − | − | + | − | − |
| ■ (light green) | + | − | + | − | − | − | − | − |
| ■ (orange) | + | − | + | − | − | − | − | + |
| ■ (red) | + | − | + | − | + | − | − | + |

Figure S2.3: Signs of local sensitivity values and their dependence on host competence and alternative host competitive ability. Within all panels, the left graph is a Voronoi diagram demonstrating how both hosts' sink and source values ($\Phi_f = \chi_f - u_f P/Y_f^*$ and $\Phi_a = \chi_a - u_a P/Y_a^*$) correspond to the signs of local sensitivity, where $\chi_f$ and $\chi_a$ are varied to obtain results. The right graph within all panels shows the values of $\chi_f$ and $\chi_a$ used to obtain the left graph, also displaying the resulting local sensitivity signs. Thus, the left graph is an alternate mapping of the data in the right graph. The table below all panels shows the signs of local sensitivity values for each colored region. Analytically, we show that $\text{sign}(\partial Y_f^*/\partial \beta_a) = -\text{sign}(\partial Y_f^*/\partial \rho_a)$, and that $\partial Y_f^*/\partial \chi_a > 0$ and $\partial Y_f^*/\partial u_a < 0$ are always true. **A, B**: A demonstration of all possible signs of local sensitivity values for a nominal parameter set by varying host shedding rates. **A** shows a broad view of parameter space, with shedding rates varying from 1 to 12. The right graph of this panel shows the inset box for the data shown in **B**. **C, D**: A demonstration of the effect of varying alternative host competitive ability ($\alpha_{fa}$) on the regions of parameter space where local sensitivity values take on different signs. Both panels show the same nominal parameter set, with **C** using $\alpha_{fa} = 0.3$ and **D** using $\alpha_{fa} = 1.2$. Other parameters for panels **A** and **B** are $b_f = 1$, $\alpha_{af} = 0.97$, $u_f = 1$, $\rho_f = 0.1$, $\mu_f = 0.1$, $b_a = 1$, $\alpha_{fa} = 0.9$, $u_a = 1$, $\rho_a = 0.3$, $\mu_a = 0.11$, $\beta_a = 1$, and $\eta_a = 0$. Other parameters for panels **C** and **D** are $b_f = 1.5$, $\alpha_{af} = 0.6$, $u_f = 1$, $\rho_f = 0.15$, $\mu_f = 0.15$, $b_a = 1$, $u_a = 1$, $\rho_a = 0.1$, $\mu_a = 0.1$, $\beta_a = 1$, and $\eta_a = 0$.

### S3 Numerical Simulation of Nominal Parameter Sets

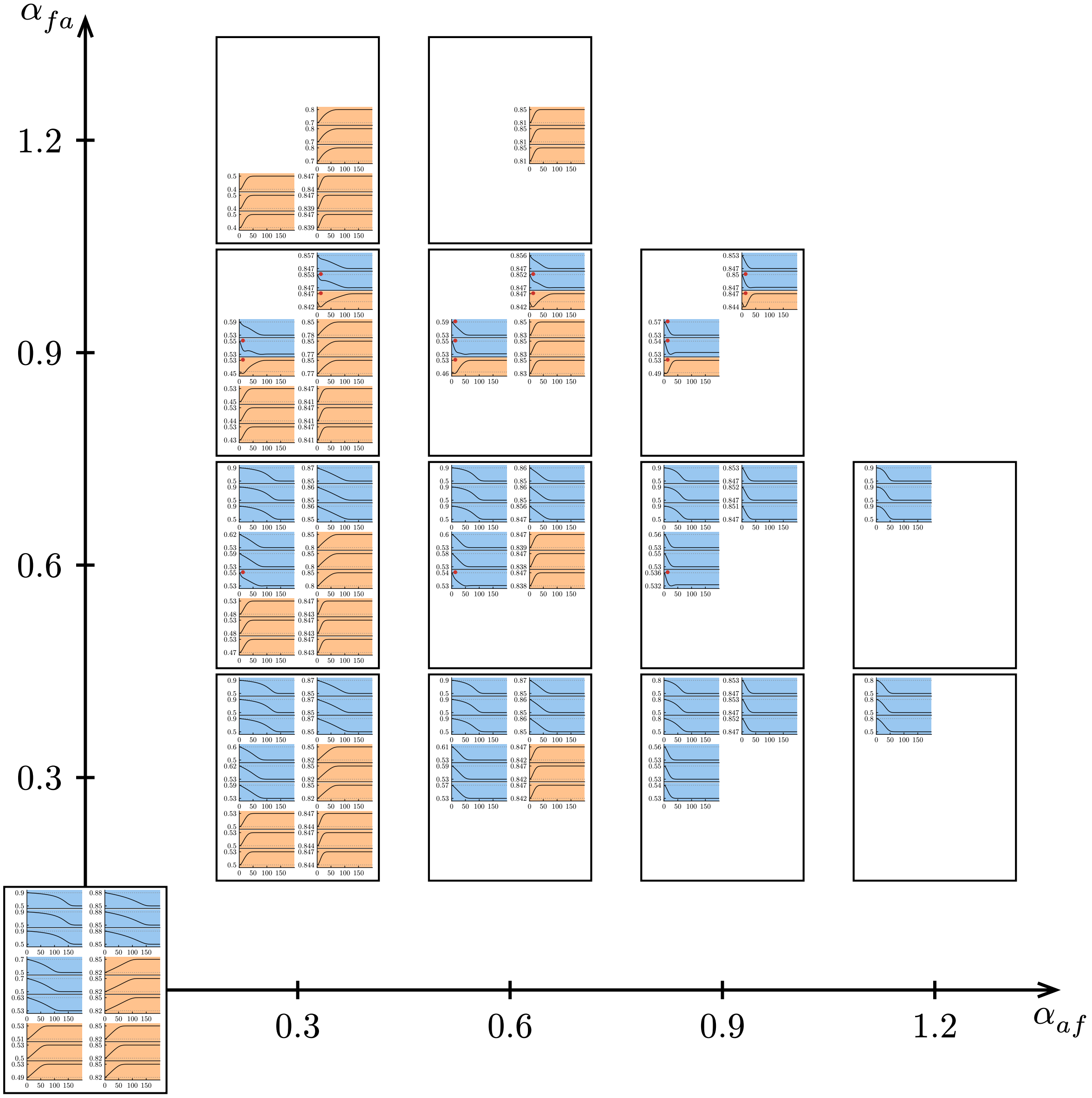

Figure S3.0: Focal host infection prevalence time series for (S1) simulation of each nominal parameter set.

### S4 Key for Global and Local Sensitivity Analysis Results

We use Partial Rank Correlation Coefficient (PRCC), first presented in McKay et al. [1976] (see Hanthanan Arachchilage et al., 2023), to calculate the global sensitivity of focal host infection prevalence to alternative host parameters. Here we give all results in a simplified form, providing the raw data separately.

| Symbol | Usage: Global Sensitivity | Usage: Local Sensitivity |
|---|---|---|
| + | positive, strongly influential (PRCC > 0.5) | positive |
| (+) | positive, moderately influential (0.1 ≤ PRCC < 0.5) | *N/A* |
| 0 | near zero, noninfluential (\|PRCC\| < 0.1) | *N/A* |
| (-) | negative, moderately influential (−0.1 ≥ PRCC > −0.5) | *N/A* |
| - | negative, strongly influential (PRCC < −0.5) | negative |

### S5 Focal Host Infection Prevalence: Simulation and Global & Local Sensitivity Analysis Results

#### S5.1 Alternative Host Shedding Rate ($\chi_a$) Effect

The local sensitivity of focal host infection prevalence ($Y_f^*$) to the alternative host shedding rate ($\chi_a$) is:

$$\frac{\partial Y_f^*}{\partial \chi_a} = -\frac{1}{\det(J)} Y_a^* N_a^* \det(J^{\cancel{53}}) \propto \left(\mu_a + \frac{b_a + \rho_a}{(1 - Y_a^*)^2}\right)(1 - \alpha_{fa}\alpha_{af}). \quad \text{(S5)}$$

The sensitivity is positive unless $\alpha_{fa}\alpha_{af} > 1$. This means that an increase in alternative host shedding rate increases focal host infection prevalence at equilibrium, unless interspecific competition is sufficiently high such that the host species cannot coexist in the absence of the pathogen ($\alpha_{fa}\alpha_{af} > 1$). For all nominal parameter sets, this sensitivity is positive because we only consider systems where the hosts can stably coexist in the absence of the pathogen (requires $\alpha_{fa}\alpha_{af} < 1$).

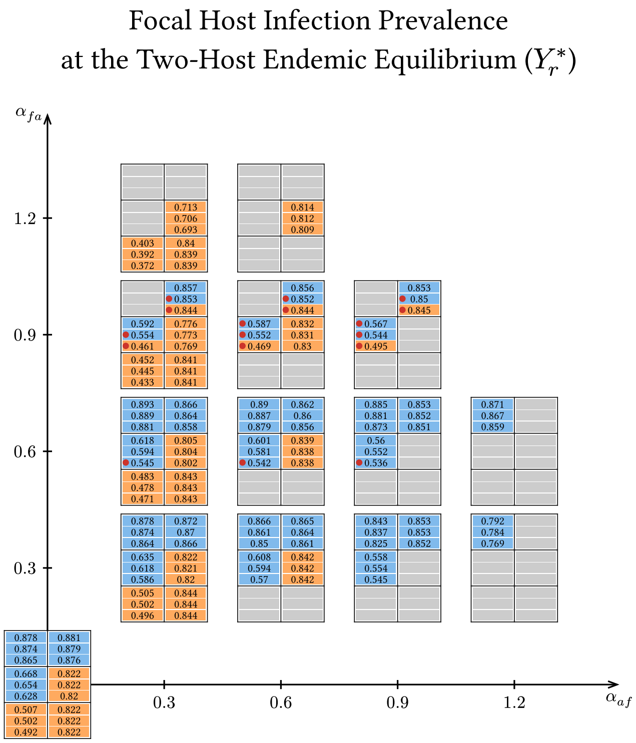

Figure S5.0: Numerical calculation of focal host infection prevalence at the two-host endemic equilibrium.

In line with local sensitivity results, global sensitivity analysis shows that higher alternative host shedding rates increase focal host infection prevalence at all time points (all positive values when |PRCC| > 0.1 in Figure SE1). These effects are strongly influential (|PRCC| ≥ 0.5) across all parameter space except when the focal host has sufficiently greater competence than the alternative host (plus signs in Figure SE1, except for (+) and zeros when $R_{0f}$ is sufficiently larger than $R_{0a}$).

#### S5.2 Alternative Host Uptake Rate ($u_a$) Effect

The local sensitivity of focal host infection prevalence ($Y_f^*$) to the alternative host uptake rate ($u_a$) is:

$$\frac{\partial Y_f^*}{\partial u_a} = \frac{1}{\det(J)} N_a^* P^* \det(J^{\not{53}}) \propto -\left(\mu_a + \frac{b_a + \rho_a}{(1 - Y_a^*)^2}\right)(1 - \alpha_{fa}\alpha_{af}). \tag{S6}$$

The sensitivity is negative unless $\alpha_{fa}\alpha_{af} > 1$. This means that an increase in alternative host uptake rate ($u_a$) decreases focal host infection prevalence at equilibrium, unless interspecific competition is sufficiently high such that the host species cannot coexist in the absence of the pathogen ($\alpha_{fa}\alpha_{af} > 1$). The conditions defining the sign of this sensitivity are exactly the opposite of those for $\chi_a$, as the two sensitivities always have the opposite sign (see (S5) and (S6)). For all nominal parameter sets, this sensitivity is negative because we only consider systems where the hosts can stably coexist in the absence of the pathogen (requires $\alpha_{fa}\alpha_{af} < 1$).

In line with local sensitivity results, global sensitivity analysis shows that higher alternative host uptake rates always decrease focal host infection prevalence at all time points (all negative values in Figure SE2). These effects are strongly influential (|PRCC| ≥ 0.5) across parameter space except when the focal host has sufficiently greater competence than the alternative host and the alternative host competes with the focal host (minus signs in Figure SE2, except for (−) when $R_{0f}$ is sufficiently larger than $R_{0a}$ and $\alpha_{fa} > 0$).

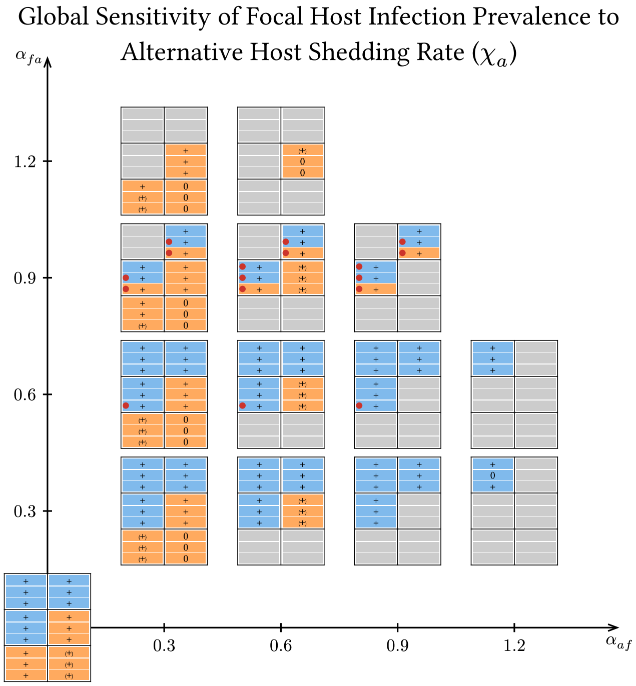

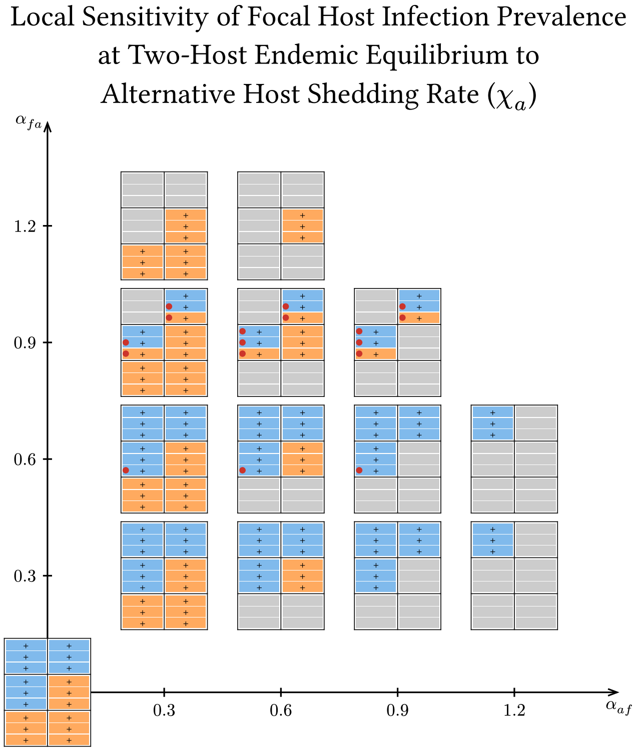

Figure S5.1: Numerical calculation of global and local sensitivity of focal host infection prevalence to alternative host shedding rate ($\chi_a$). Global sensitivity results take multiple timesteps into account, whereas local sensitivity only evaluates sensitivity at the two-host endemic equilibrium.

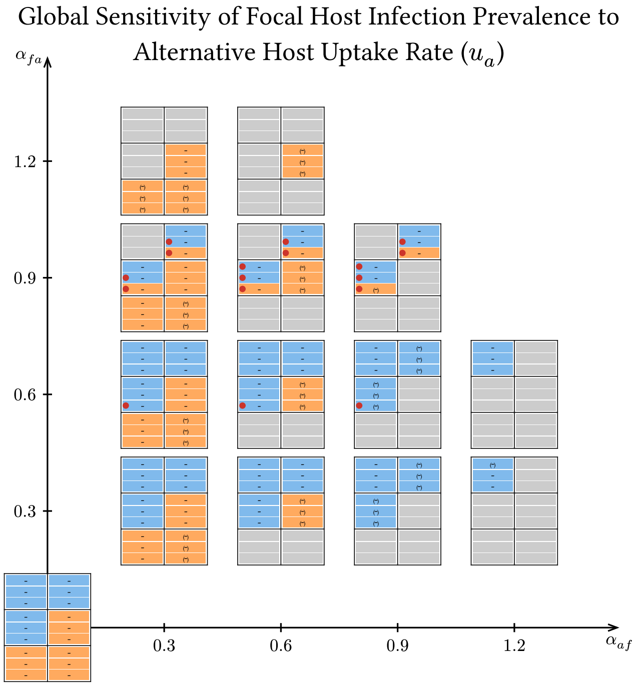

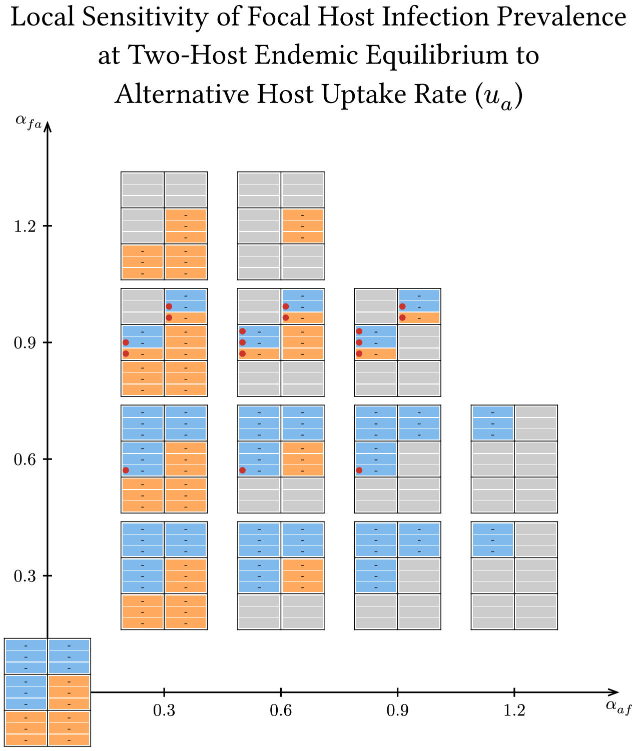

Figure S5.2: Numerical calculation of global and local sensitivity of focal host infection prevalence to alternative host uptake rate ($u_a$). Global sensitivity results take multiple timesteps into account, whereas local sensitivity only evaluates sensitivity at the two-host endemic equilibrium.

#### S5.3 Alternative Host Infection Rate ($\beta_a$) Effect

The local sensitivity of focal host infection prevalence ($Y_f^*$) to the alternative host infection rate ($\beta_a$) is

$$\frac{\partial Y_f^*}{\partial \beta_a} = \frac{1}{\det(J)} P^* \det(J^{\not{43}}) \propto (1 - \alpha_{fa}\alpha_{af})\chi_a N_a^* + \mu_a(\alpha_{fa} Y_f^* \Phi_f - Y_a^* \Phi_a). \tag{S7}$$

The sensitivity is positive in much of parameter space, meaning that focal host infection prevalence at equilibrium is higher when the alternative host has a higher infection rate. However, the sensitivity can be negative when (i) interspecific competition is sufficiently strong such that the pathogen mediates coexistence of the hosts ($\alpha_{fa}\alpha_{af} > 1$), (ii) the alternative host is a sufficiently large source ($\Phi_a$ positive and large), or (iii) the alternative host is a sufficiently strong interspecific competitor and the focal host is a sufficiently large sink ($\alpha_{fa}$ large and $\Phi_f$ negative and large).

Global sensitivity analysis shows that higher alternative host infection rates ($\beta_a$) increase focal host infection prevalence at all time points (all positive values when |PRCC| > 0.1 in Figure SE3). These effects are strongly influential (|PRCC| ≥ 0.5) when the competence of the alternative host is similar to or greater than the focal host's (plus signs only occur in Figure SE3 when $R_{0a}$ is greater than or similar to $R_{0f}$). However, the effects are only moderately influential (0.1 ≤ |PRCC| < 0.5) or noninfluential (|PRCC| < 0.1) when the competence of the alternative host is much less than the focal host's ((+) and zeros in Figure SE3 when $R_{0a}$ is much less than $R_{0f}$). Local sensitivity analysis suggests that nominal parameter sets in more extreme regions of parameter space could show the opposite effect, where higher infection rates decrease focal host infection prevalence (see Figure SB4-2).

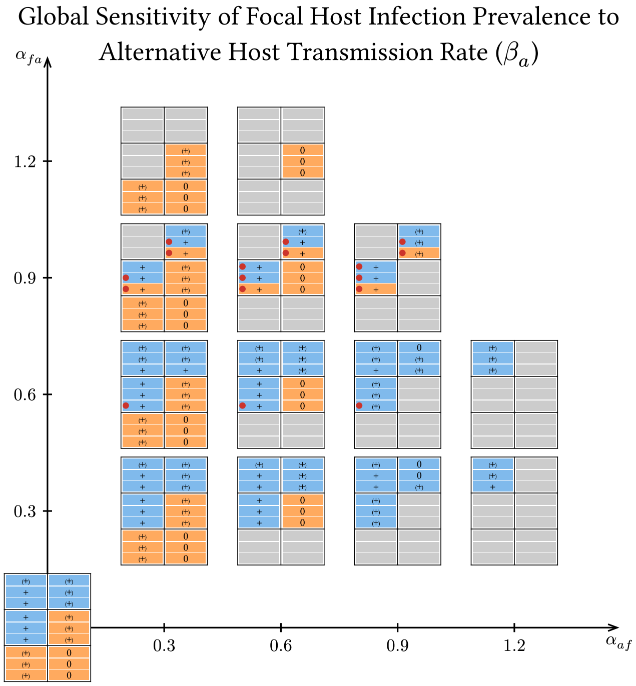

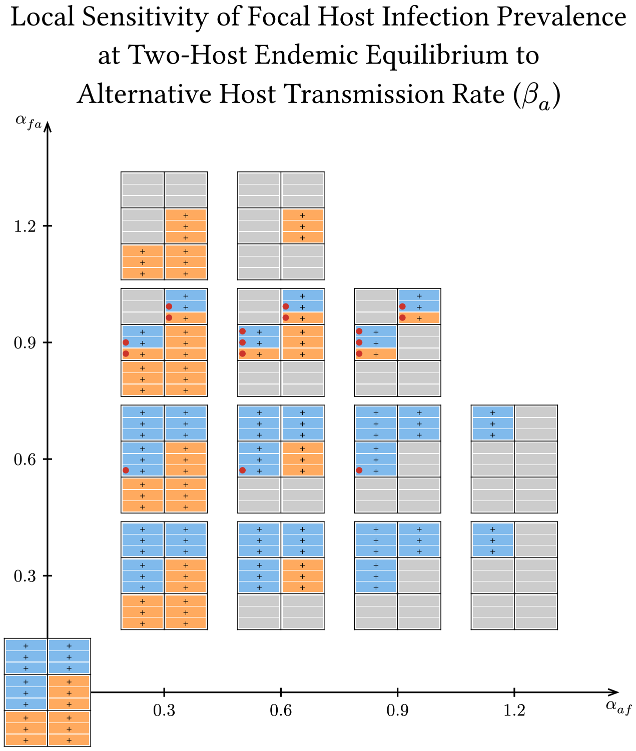

Figure S5.3: Numerical calculation of global and local sensitivity of focal host infection prevalence to alternative host infection rate ($\beta_a$). Global sensitivity results take multiple timesteps into account, whereas local sensitivity only evaluates sensitivity at the two-host endemic equilibrium.

#### S5.4 Alternative Host Recovery Rate ($\rho_a$) Effect

The local sensitivity of focal host infection prevalence ($Y_f^*$) to the alternative host recovery rate ($\rho_a$) is:

$$\frac{\partial Y_f^*}{\partial \rho_a} = -\frac{1}{\det(J)} \frac{Y_a^*}{1 - Y_a^*} \det(J^{\not{4}\not{3}}) \propto -(1 - \alpha_{fa}\alpha_{af})\chi_a N_a^* - \mu_a(\alpha_{fa} Y_f^* \Phi_f - Y_a^* \Phi_a). \tag{S8}$$

The sensitivity is negative in much of parameter space, meaning that focal host infection prevalence at equilibrium is lower when the alternative host has a higher recovery rate. However, the sensitivity can be positive when (i) interspecific competition is sufficiently strong such that the pathogen mediates coexistence of the hosts ($\alpha_{fa}\alpha_{af} > 1$), (ii) the alternative host is a sufficiently large source ($\Phi_a$ positive and large), or (iii) the alternative host is a sufficiently strong interspecific competitor and the focal host is a sufficiently large sink ($\alpha_{fa}$ large and $\Phi_f$ negative and large). These conditions defining exceptions are identical to those for $\beta_a$, as the two sensitivities always have the opposite sign (see (S7) and (S8)).

Global sensitivity analysis shows that higher alternative host recovery rates ($\rho_a$) decrease focal host infection prevalence at all time points (negative values when $|\text{PRCC}| > 0.1$ in Figure SE4). These effects are never strongly influential ($|\text{PRCC}| \geq 0.5$) (no minus signs in Figure SE4). They are only moderately influential ($0.1 \leq |\text{PRCC}| < 0.5$) when hosts have similar competences ((−) only occur when $R_{0f}$ is similar to $R_{0a}$ in Figure SE4), and are always noninfluential ($|\text{PRCC}| < 0.1$) when hosts have dissimilar competences (all zeros when $R_{0f}$ is much greater or much less than $R_{0a}$ in Figure SE4). Local sensitivity analysis suggests that nominal parameter sets in more extreme regions of parameter space could show the opposite effect, where higher infection rates decrease focal host infection prevalence (see Figure SB4-2).

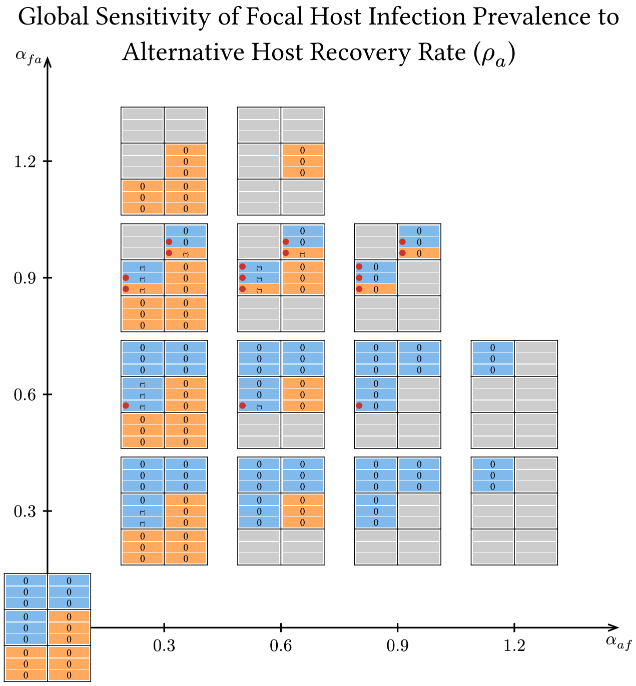

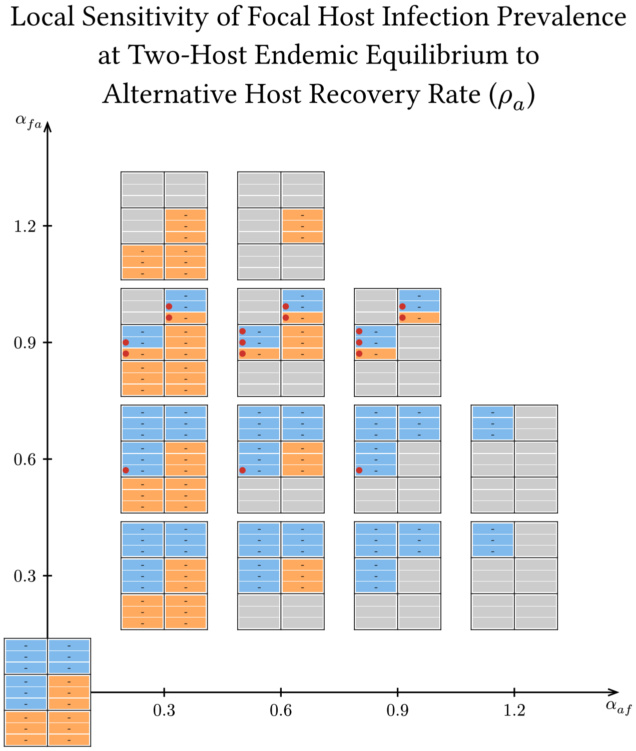

Figure S5.4: Numerical calculation of global and local sensitivity of focal host infection prevalence to alternative host recovery rate ($\rho_a$). Global sensitivity results take multiple timesteps into account, whereas local sensitivity only evaluates sensitivity at the two-host endemic equilibrium.

#### S5.5 Alternative Host Disease Mortality Rate ($\mu_a$) Effect

The local sensitivity of focal host infection prevalence ($Y_f^*$) to the alternative host disease mortality rate ($\mu_a$) is:

$$\begin{aligned}\frac{\partial Y_f^*}{\partial \mu_a} &= -\frac{1}{\det(J)} Y_a^* \left(\det(J^{\not{2}\not{3}}) + \det(J^{\not{4}\not{3}})\right) \\ &\propto -(1 - \alpha_{fa}\alpha_{af})\chi_a N_a^* + \frac{b_a + \rho_a}{1 - Y_a^*}(\alpha_{fa} Y_f^* \Phi_f - Y_a^* \Phi_a).\end{aligned}$$

The sensitivity is negative in much of parameter space, meaning that focal host infection prevalence at equilibrium is lower when the alternative host has a higher disease mortality rate. However, the sensitivity can be positive when (i) interspecific competition is sufficiently strong such that the pathogen mediates coexistence of the hosts ($\alpha_{fa}\alpha_{af} > 1$), (ii) the alternative host is a sufficiently large sink ($\Phi_a$ negative and large), or (iii) the alternative host is a sufficiently strong interspecific competitor and the focal host is a sufficiently large source ($\alpha_{fa}$ large and $\Phi_f$ positive and large).

Global sensitivity analysis shows that higher alternative host disease mortality rates ($\mu_a$) decrease focal host infection prevalence at all time points when the competence of the alternative host is greater than or similar to the focal host's (all negative values when $R_{0a}$ is greater than or similar to $R_{0f}$ in Figure SE5), and increase it when the competence of the alternative host is much less than the focal host's (all positive values when $R_{0a}$ is much less than $R_{0f}$ in Figure SE5). These effects are only strongly influential ($|\text{PRCC}| \geq 0.5$) when the competence of the alternative host is much greater than the competence of the focal host and the focal host is a strong competitor (minus signs only occur when $R_{0a}$ is much greater than $R_{0f}$ and $\alpha_{af} = 1.2$ in Figure SE5). In all other areas of parameter space, the parameter is moderately influential ($0.1 \leq |\text{PRCC}| < 0.5$) or noninfluential ($|\text{PRCC}| < 0.1$) ((+), (−), and 0 when $\alpha_{af} \neq 1.2$ in Figure SE5).

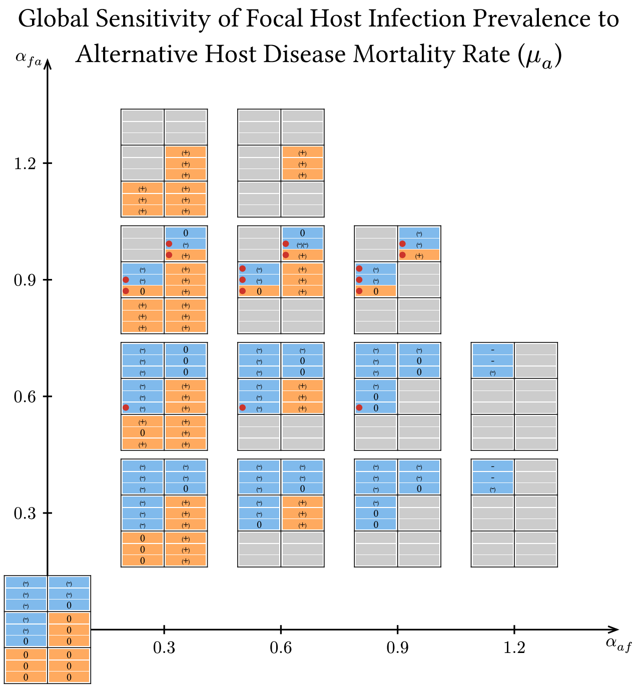

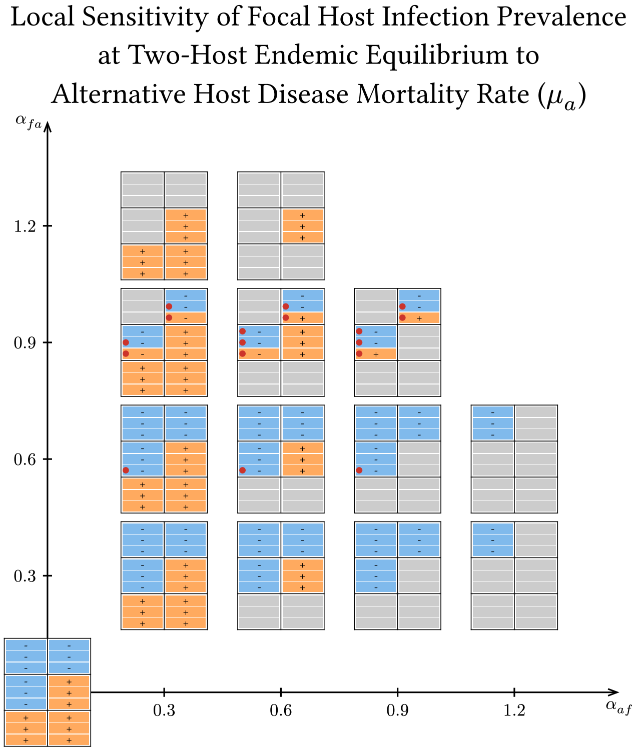

Figure S5.5: Numerical calculation of global and local sensitivity of focal host infection prevalence to alternative host disease mortality rate ($\mu_a$). Global sensitivity results take multiple timesteps into account, whereas local sensitivity only evaluates sensitivity at the two-host endemic equilibrium.

#### S5.6 Alternative Host Birth Rate ($b_a$) Effect

The local sensitivity of focal host infection prevalence ($Y_f^*$) to the alternative host birth rate ($b_a$) is:

$$\begin{aligned}\frac{\partial Y_f^*}{\partial b_a} &= \frac{1}{\det(J)}\left(\det(J^{\cancel{23}}) - \frac{Y_a^*}{1-Y_a^*}\det(J^{\cancel{43}})\right)\\ &\propto -Y_a^*(1-\alpha_{fa}\alpha_{af})\chi_a N_a^* - \left(\mu_a + \frac{b_a+\rho_a}{1-Y_a^*}\right)(\alpha_{fa}Y_f^*\Phi_f - Y_a^*\Phi_a).\end{aligned}$$

The sensitivity is negative in much of parameter space, meaning that focal host infection prevalence at equilibrium is lower when the alternative host has a higher birth rate. However, the sensitivity can be positive when (i) interspecific competition is sufficiently strong such that the pathogen mediates coexistence of the hosts ($\alpha_{fa}\alpha_{af} > 1$), (ii) the alternative host is a sufficiently large source ($\Phi_a$ positive and large), or (iii) the alternative host is a sufficiently strong interspecific competitor and the focal host is a sufficiently large sink ($\alpha_{fa}$ large and $\Phi_f$ negative and large).

Global sensitivity analysis shows that higher alternative host birth rates ($b_a$) almost always increase focal host infection prevalence at all time points when the competence of the alternative host is greater than or similar to the focal host's (almost always positive when $R_{0a}$ is greater than or similar to $R_{0f}$ when $|\text{PRCC}| > 0.1$ in Figure SE6), and decrease it when the competence of the alternative host is much less than the focal host's (all negative signs when $R_{0a}$ is much less than $R_{0f}$ in Figure SE6). Exceptions to this pattern only occur at or near non-monotonic cases, where higher alternative host birth rates decrease focal host infection prevalence despite hosts having similar competence (deviations from the pattern only occur with negative values when $R_{0a}$ is similar to $R_{0f}$ and at or near red dot cases in Figure SE6) and in one instance where higher alternative host birth rates decrease focal host infection prevalence initially, but later increase it (one cell with (−)(+) near red dot cases at $\alpha_{af} = 0.3$, $\alpha_{fa} = 0.9$). These effects are strongly influential ($|\text{PRCC}| \geq 0.5$) except when host species have similar competence, where a broad range of influence is observed across cases (plus and minus signs except when $R_{0a}$ is similar to $R_{0f}$, which have plus, minus, (+), (−), and zero in Figure SE6). The case where increasing $b_a$ had a different direction of effect at different time scales was only moderately influential ($0.1 \leq |\text{PRCC}| < 0.5$).

#### S5.7 Alternative Host Competitive Ability ($\alpha_{fa}$) Effect

The local sensitivity of focal host infection prevalence ($Y_f^*$) to the alternative host competition parameter ($\alpha_{fa}$) is:

$$\frac{\partial Y_f^*}{\partial \alpha_{fa}} = \frac{1}{\det(J)} N_a^* \det(J^{\not 13}) \propto \alpha_{af} Y_a^* \Phi_a - Y_f^* \Phi_f.$$

The sensitivity can take on either sign, meaning that increased competitive ability of the alternative host increases or decreases focal host infection prevalence when the focal host is a sink ($\Phi_f < 0$) or source ($\Phi_f > 0$), respectively. However, if the focal host is a sufficiently strong interspecific competitor ($\alpha_{af}$ large), then the response can be reversed when the alternative host is a sufficiently large sink ($\Phi_a < 0$) or source ($\Phi_a > 0$), respectively.

Global sensitivity analysis shows that higher alternative host competitive abilities ($\alpha_{fa}$) decrease focal host infection prevalence at all time points when the alternative host competence is similar to or lower than the focal host's (minus signs when $R_{0a}$ is similar to or less than $R_{0f}$ in Figure SE7), and increase focal host infection prevalence otherwise (plus signs when $R_{0a}$ is much greater than $R_{0f}$ in Figure SE7). When the alternative host is a stronger competitor than the focal host, this effect is strongly influential ($|\text{PRCC}| \geq 0.5$) only when the alternative host does not have low competence or the focal host does not have high competence (when $\alpha_{fa} > \alpha_{af}$, minus signs and plus signs never occur in the bottom right block of three cells and occur at least once in every other block in Figure SE7). When the alternative host is a weaker competitor than the focal host, this effect is strongly influential ($|\text{PRCC}| \geq 0.5$) only when the competence of the alternative host

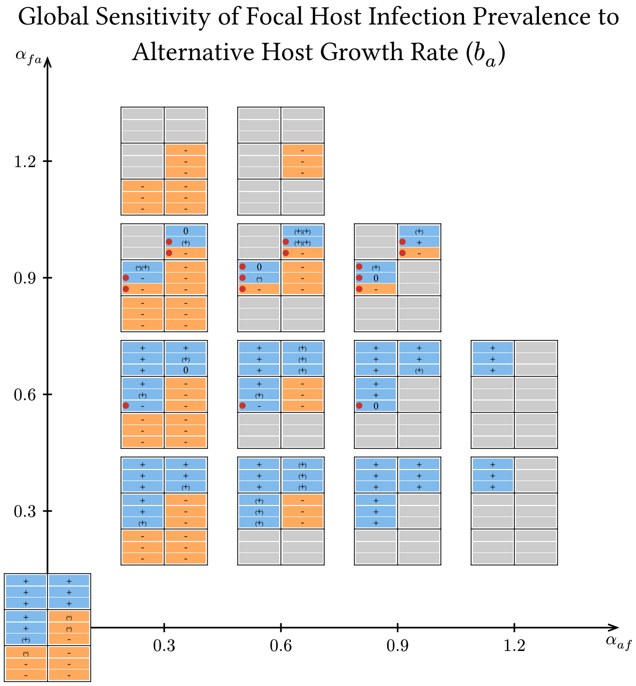

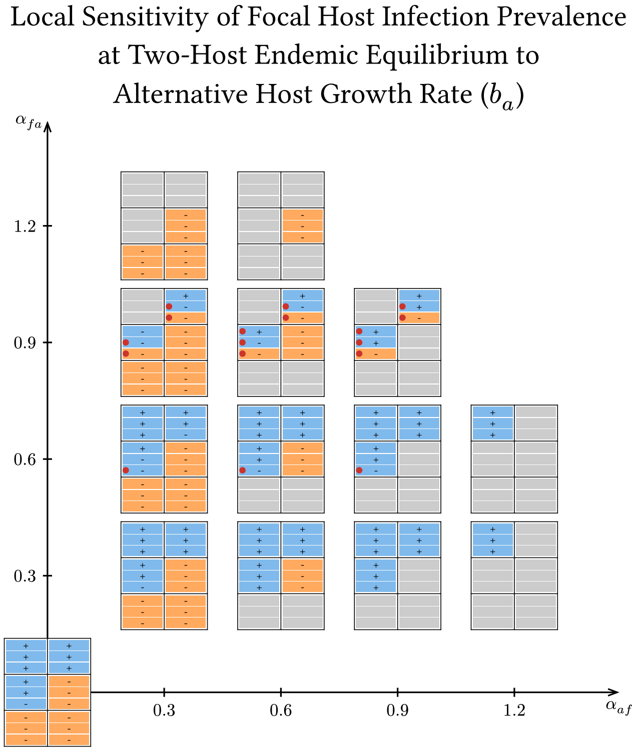

Figure S5.6: Numerical calculation of global and local sensitivity of focal host infection prevalence to alternative host birth rate ($b_a$). Global sensitivity results take multiple timesteps into account, whereas local sensitivity only evaluates sensitivity at the two-host endemic equilibrium.

is much greater than the focal host's (when $\alpha_{fa} < \alpha_{af}$, no minus signs occur, and plus signs only occur in the top left block of three cells in Figure SE7). When hosts have equal competitive ability, this effect is strongly influential ($|\text{PRCC}| \geq 0.5$) only when at least one host has moderate competence and the alternative host does not have low competence (when $\alpha_{fa} = \alpha_{af}$, minus signs and plus signs only occur in the top left, middle left, and middle right blocks in Figure SE7).

#### S5.8 Local Sensitivity to Alternative Host Loss Rate ($\eta_a$)

To assess how loss of the alternative host affects focal host infection prevalence over time when the loss rate is very slow, we replace the time-dependent loss rate ($\zeta_a t$) in (**??**) with a time-independent loss rate ($\eta_a$) and evaluate the sensitivity of focal host infection prevalence at equilibrium to $\eta_a$. In other words, the sign in the sensitivity of focal host infection prevalence at equilibrium ($Y_f^*$) to $\eta_a$ indicates the initial direction of change in focal host infection prevalence when $\zeta_a$ is sufficiently small.

The local sensitivity of focal host infection prevalence ($Y_f^*$) to the alternative host loss rate ($\eta_a$) is

$$\frac{\partial Y_f^*}{\partial \eta_a} = -\frac{1}{\det(J)} \det(J^{\not{2}\not{3}}) \propto \alpha_{fa} Y_f^* \Phi_f - Y_a^* \Phi_a.$$

The sensitivity can take on either sign, meaning that an increased time-independent loss rate of the alternative host increases or decreases focal host infection prevalence when the alternative host is a sink ($\Phi_a < 0$) or source ($\Phi_a > 0$), respectively. However, if the alternative host is a sufficiently strong interspecific competitor ($\alpha_{fa}$ large), then the response can be reversed when the focal host is a sufficiently large source ($\Phi_f > 0$) or sink ($\Phi_f < 0$), respectively.

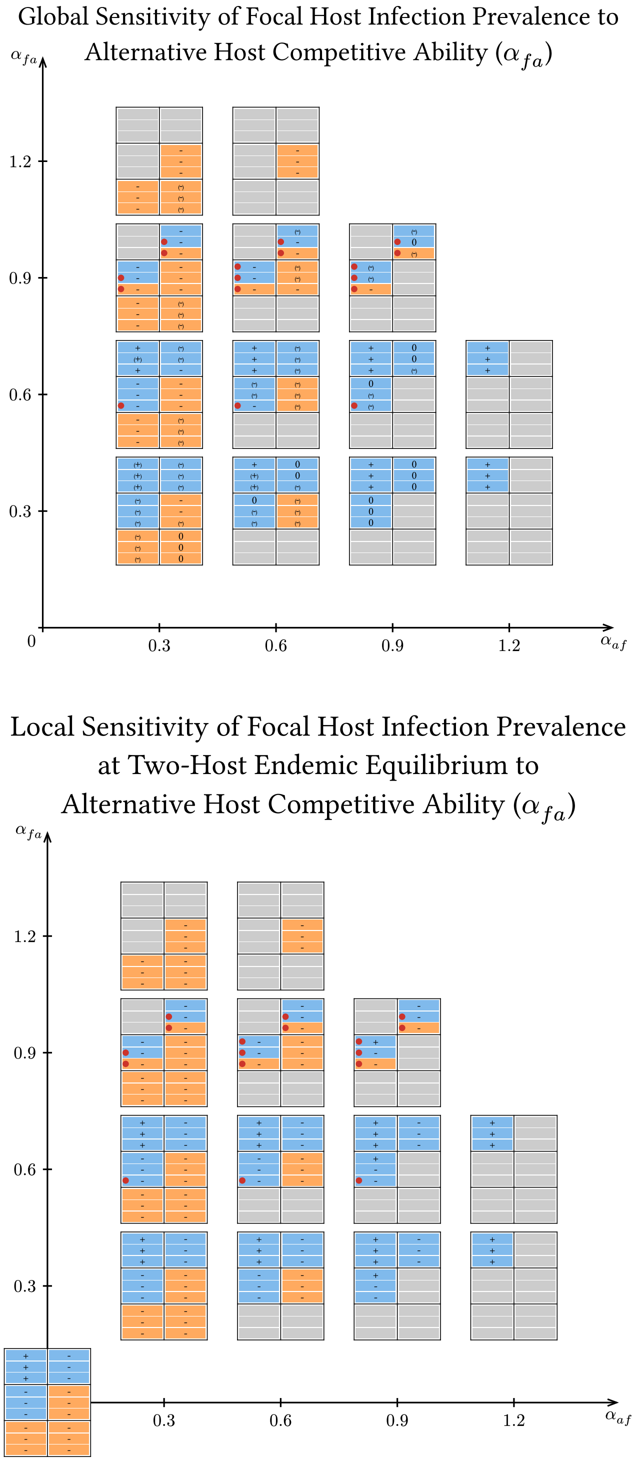

Figure S5.7: Numerical calculation of global and local sensitivity of focal host infection prevalence to alternative host competitive ability ($\alpha_{fa}$). Global sensitivity results take multiple timesteps into account, whereas local sensitivity only evaluates sensitivity at the two-host endemic equilibrium.

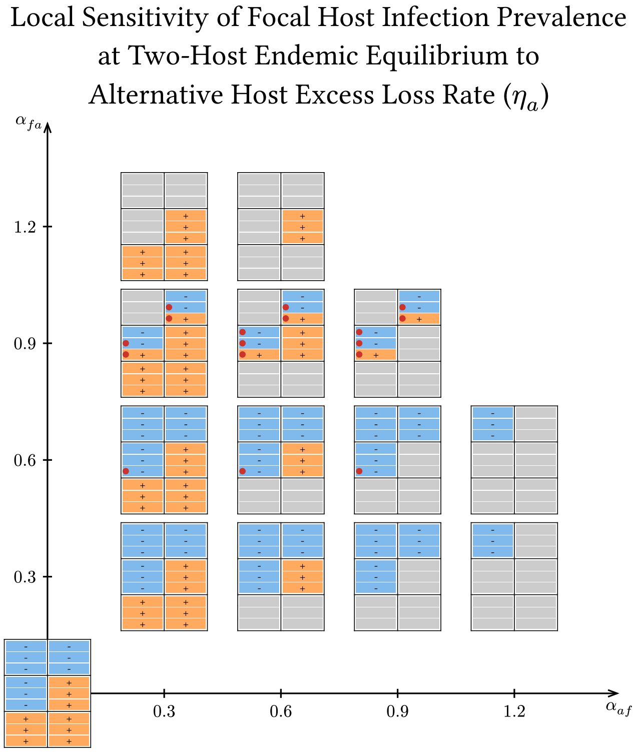

Figure S5.8: Numerical calculation of local sensitivity of focal host infection prevalence to alternative host loss rate ($\eta_a$), which we used in place of $\zeta_a t$ in this analysis. These results are equivalent to the direction that focal host infection prevalence would change at the onset of alternative host loss if alternative host loss parameter $\zeta_a$ is very small.

### S6 Divergence Time: Simulation and Global Sensitivity Analysis Results

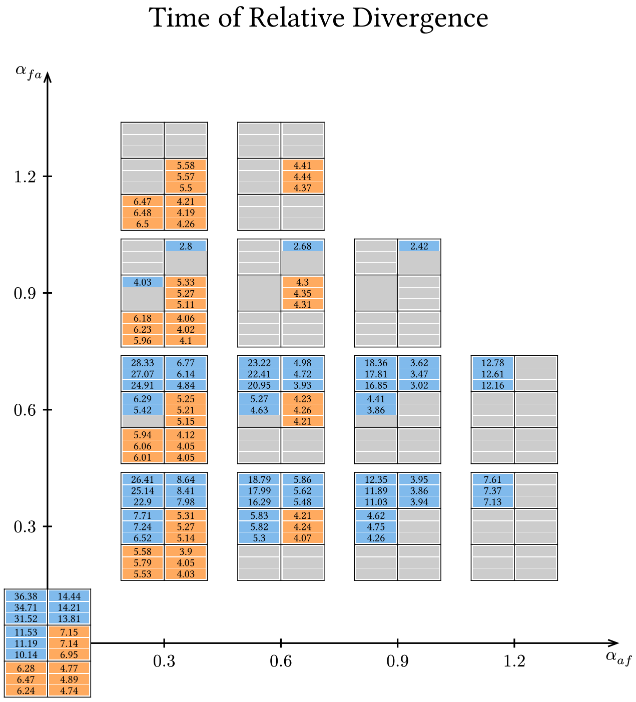

Figure S6.0: Numerical calculation of the time of relative divergence at the two-host endemic equilibrium.

#### S6.1 Alternative Host Shedding Rate ($\chi_a$) Effect on Divergence Time

Higher alternative host shedding rates ($\chi_a$) increase divergence time (all positive when $|\text{PRCC}| > 0.1$ in Figure SF1). This effect is strongly influential ($|\text{PRCC}| \geq 0.5$) when the competence of the alternative host is sufficiently large compared to the focal host's (plus signs only occur outside of the bottom right block of three cells in Figure SF1).

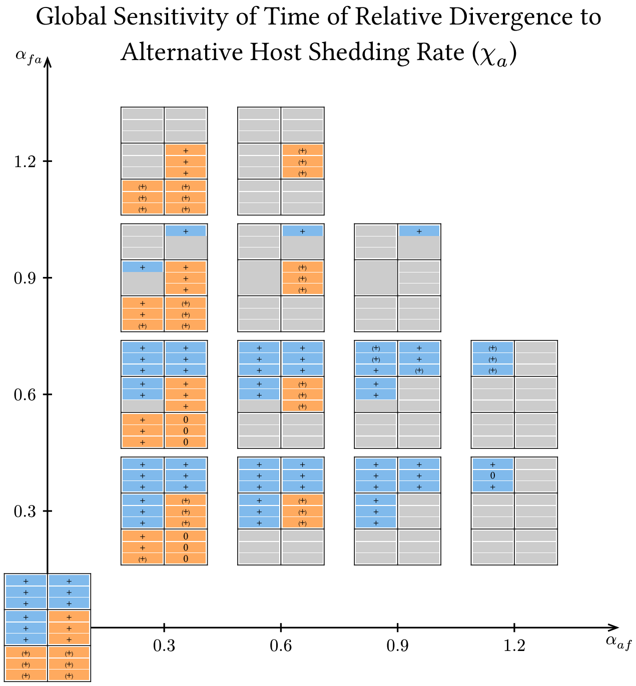

Figure S6.1: Global sensitivity of the time of relative divergence to alternative host shedding rate ($\chi_a$).

#### S6.2 Alternative Host Uptake Rate ($u_a$) Effect on Divergence Time

Higher alternative host uptake rates ($u_a$) nearly always decrease divergence time (negative nearly everywhere when |PRCC| > 0.1 in Figure SF2). Exceptions occur when hosts do not compete against each other ($\alpha_{af} = \alpha_{fa} = 0$) and when both hosts have moderate or low competence, where it increases divergence time (positive in bottom left and middle left blocks when |PRCC| > 0.1 for grid at the origin in Figure SF2). When hosts compete with each other, the effect is only strongly influential (|PRCC| ≥ 0.5) when the competence of the alternative host is less than or similar to the focal host's (plus signs only occur when $R_{0a}$ is less than or similar to $R_{0f}$, in Figure SF2). When hosts do not compete with each other, the effect is never strongly influential (|PRCC| ≥ 0.5) ((−), (+), and zeros in grid at the origin in Figure SF2).

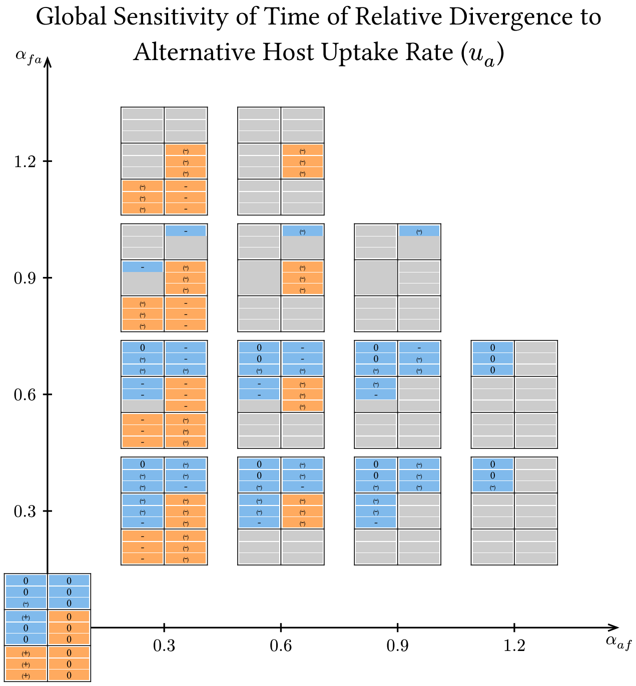

Figure S6.2: Global sensitivity of the time of relative divergence to alternative host uptake rate ($u_a$).

#### S6.3 Alternative Host Infection Rate ($\beta_a$) Effect on Divergence Time

Higher alternative host infection rates ($\beta_a$) increase divergence time (all plus signs when |PRCC| > 0.1 in Figure SF3). This effect is only strongly influential (|PRCC| ≥ 0.5) when focal host competence is moderate and alternative host competence is moderate or high (only plus signs occur in top left and middle left blocks of three cells across grids in Figure SF3).

#### S6.4 Alternative Host Recovery Rate ($\rho_a$) Effect on Divergence Time

Higher alternative host recovery rates ($\rho_a$) decrease divergence time (negative when |PRCC| > 0.1 in Figure SF4). However, this effect is noninfluential (|PRCC| < 0.1) except for two

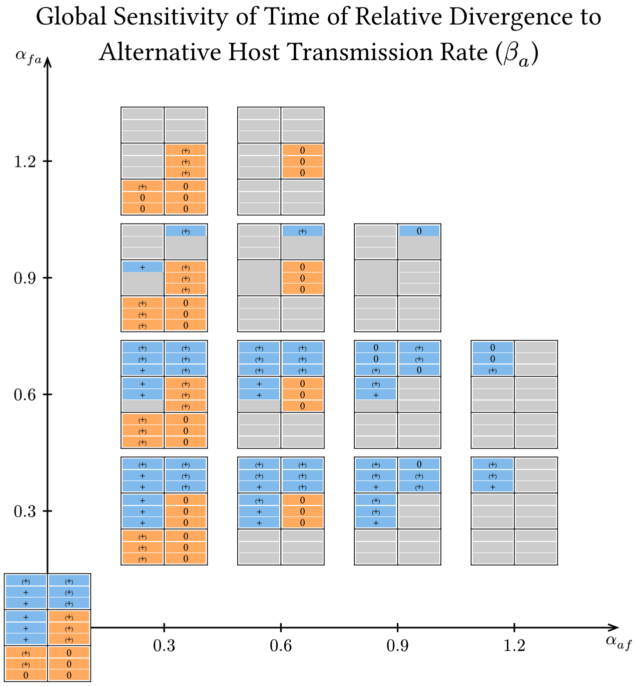

Figure S6.3: Global sensitivity of the time of relative divergence to alternative host infection rate ($\beta_a$).

cases, where it is moderately influential ($0.1 \leq |\text{PRCC}| < 0.5$) (two (−) and otherwise zeros in Figure SF4).

#### S6.5 Alternative Host Disease Mortality Rate ($\mu_a$) Effect on Divergence Time

Higher alternative host disease mortality rates ($\mu_a$) decrease divergence time (negative when $|\text{PRCC}| > 0.1$ in Figure SF5). This effect is almost always moderately influential ($0.1 \leq |\text{PRCC}| < 0.5$), and is only strongly influential ($|\text{PRCC}| \geq 0.5$) when the focal host is a strong competitor and alternative host competence is much higher than focal host competence ((−) and some zeros besides when $\alpha_{af} = 1.2$, which has minus signs, in Figure SF5).

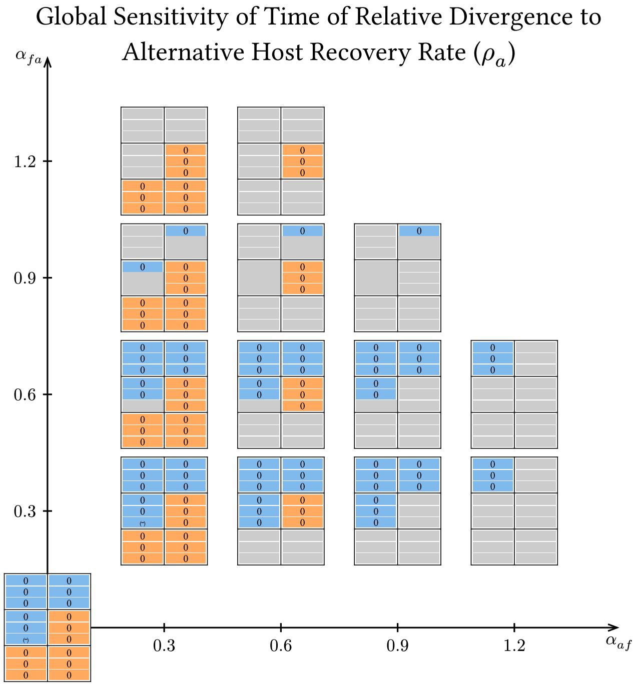

Figure S6.4: Global sensitivity of the time of relative divergence to alternative host recovery rate ($\rho_a$).

#### S6.6 Alternative Host Birth Rate ($b_a$) Effect on Divergence Time

Higher alternative host birth rates ($b_a$) nearly always increase divergence time (positive nearly everywhere when $|\text{PRCC}| > 0.1$ in Figure SF6). One exception occurs when the focal host is a weak competitor, the alternative host is a moderately strong competitor, and both hosts have moderate competence, where it decreases divergence time (one negative when $\alpha_{af} = 0.3$, $\alpha_{fa} = 0.9$, and $R_{0a}$ and $R_{0f}$ are moderate in Figure SF6). The effect of this parameter is strongly influential ($|\text{PRCC}| \geq 0.5$) everywhere except when host competences are similar (plus signs everywhere when $R_{0a}$ and $R_{0f}$ are dissimilar in Figure SF6), where the full range of influences is observed (plus signs, (+), and zeros when $R_{0a}$ and $R_{0f}$ are similar in Figure SF6). The exceptional case where higher $b_a$ decreases divergence time is moderately influential ($0.1 \leq |\text{PRCC}| < 0.5$).

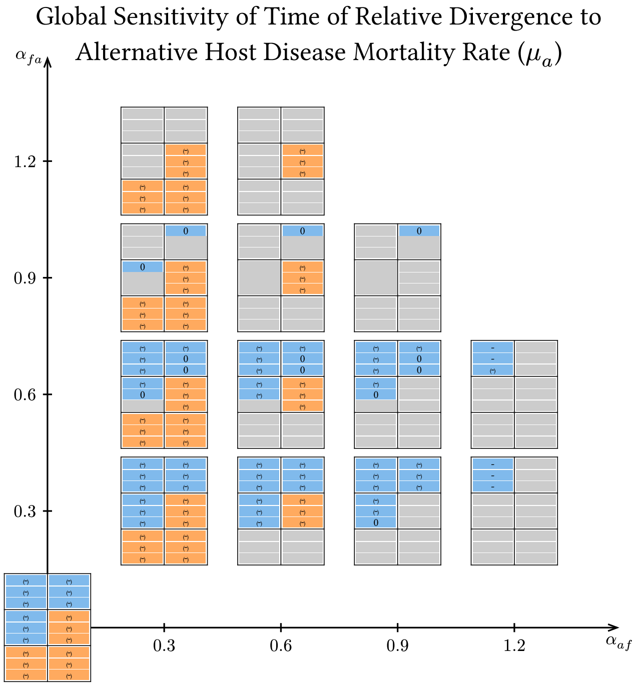

Figure S6.5: Global sensitivity of the time of relative divergence to alternative host disease mortality rate ($\mu_a$).

#### S6.7 Alternative Host Competitive Ability ($\alpha_{fa}$) Effect on Divergence Time

Higher alternative host competitive ability ($\alpha_{fa}$) increases divergence time when hosts have sufficiently different competences (positive when $R_{0a}$ is dissimilar to $R_{0f}$ and when |PRCC| > 0.1 in Figure SF7), while it decreases divergence time when hosts have similar competences (negative when $R_{0a}$ is similar to $R_{0f}$ in Figure SF7). When the alternative host is a weak competitor, this effect is only strongly influential (|PRCC| ≥ 0.5) when the alternative host has low competence and the focal host has moderate competence (when $\alpha_{fa}$ = 0.3, grids only have plus signs or minus signs in top left or middle left blocks in Figure SF7). When the alternative host is a moderately weak competitor, this effect is only strongly influential (|PRCC| ≥ 0.5) when the alternative host has high competence or the focal host has moderate competence (when $\alpha_{fa}$ = 0.6, grids only have plus signs

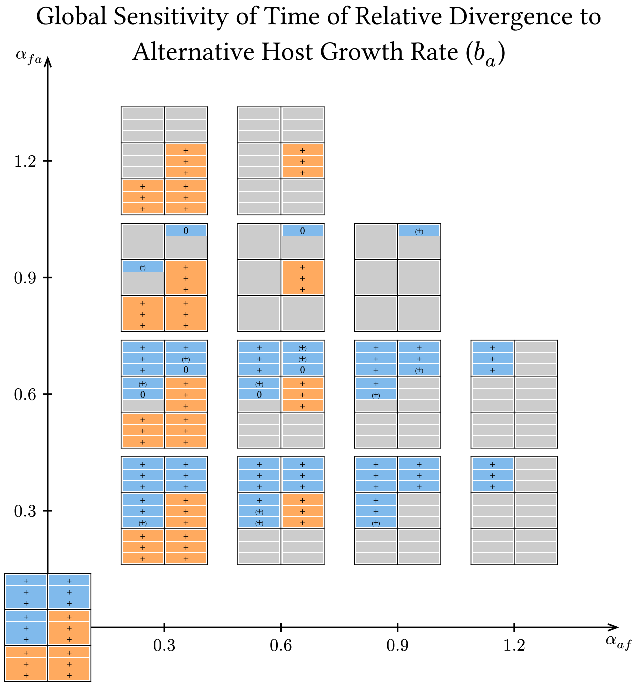

Figure S6.6: Global sensitivity of the time of relative divergence to alternative host birth rate ($b_a$).

or minus signs outside of bottom right and middle right blocks in Figure SF7). When the alternative host is a moderately strong competitor, this effect is only strongly influential (|PRCC| ≥ 0.5) when the alternative host has low competence or when both hosts have moderate competence (when $\alpha_{fa}$ = 0.9, grids only have plus signs or minus signs in bottom right, middle right, and middle left blocks in Figure SF7). Finally, when the alternative host is a strong competitor, this effect is only strongly influential (|PRCC| ≥ 0.5) when the competence of the alternative host is low or the focal host competence is high, but not both (when $\alpha_{fa}$ = 1.2, grids only have plus signs or minus signs in bottom right, middle right, and middle left blocks in Figure SF7).

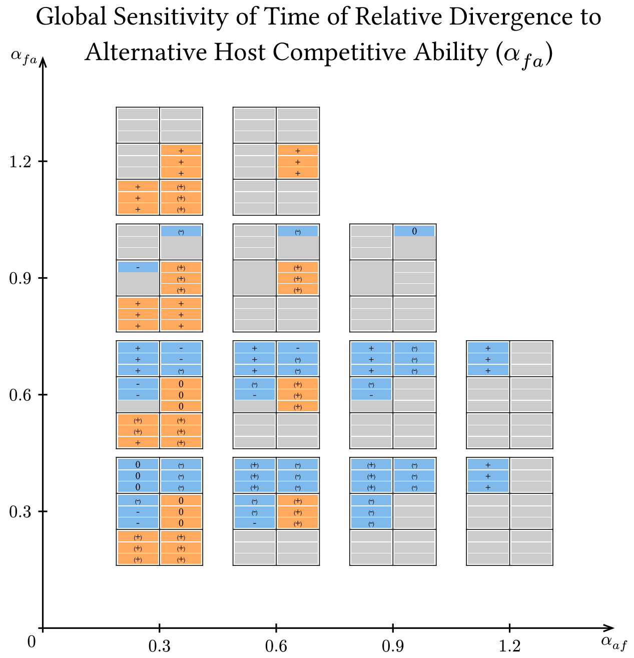

Figure S6.7: Global sensitivity of the time of relative divergence to alternative host competitive ability ($\alpha_{fa}$).

M McKay, W Conover, and D Whiteman. Report on the application of statistical techniques to the analysis of computer codes, 08 1976.

Peter Yodzis. The indeterminacy of ecological interactions as perceived through perturbation experiments. *Ecology*, 69(2):508–515, 4 1988. doi: 10.2307/1940449.
